# Population heterogeneity in the epithelial to mesenchymal transition is controlled by NFAT and phosphorylated Sp1

**DOI:** 10.1101/028993

**Authors:** Russell Gould, David M. Bassen, Anirikh Chakrabarti, Jeffrey D. Varner, Jonathan Butcher

## Abstract

Epithelial to mesenchymal transition (EMT) is an essential differentiation program during tissue morphogenesis and remodeling. EMT is induced by soluble transforming growth factor *β* (TGF-*β*) family members, and restricted by vascular endothelial growth factor family members. While many downstream molecular regulators of EMT have been identified, these have been largely evaluated individually without considering potential crosstalk. In this study, we created an ensemble of dynamic mathematical models describing TGF-*β* induced EMT to better understand the operational hierarchy of this complex molecular program. We used ordinary differential equations (ODEs) to describe the transcriptional and post-translational regulatory events driving EMT. Model parameters were estimated from multiple data sets using multiobjective optimization, in combination with cross-validation. TGF-*β* exposure drove the model population toward a mesenchymal phenotype, while an epithelial phenotype was enhanced following vascular endothelial growth factor A (VEGF-A) exposure. Simulations predicted that the transcription factors phosphorylated SP1 and NFAT were master regulators promoting or inhibiting EMT, respectively. Surprisingly, simulations also predicted that a cellular population could exhibit phenotypic heterogeneity (characterized by a significant fraction of the population with both high epithelial and mesenchymal marker expression) if treated simultaneously with TGF-*β* and VEGF-A. We tested this prediction experimentally in both MCF10A and DLD1 cells and found that upwards of 45% of the cellular population acquired this hybrid state in the presence of both TGF-*β* and VEGF-A. We experimentally validated the predicted NFAT/Sp1 signaling axis for each phenotype response. Lastly, we found that cells in the hybrid state had significantly different functional behavior when compared to VEGF-A or TGF-*β* treatment alone. Together, these results establish a predictive mechanistic model of EMT susceptibility, and potentially reveal a novel signaling axis which regulates carcinoma progression through an EMT versus tubulogenesis response.

**Author Summary:** Tissue formation and remodeling requires a complex and dynamic balance of interactions between epithelial cells, which reside on the surface, and mesenchymal cells that reside in the tissue interior. During embryonic development, wound healing, and cancer, epithelial cells transform into a mesenchymal cell to form new types of tissues. It is important to understand this process so that it can be controlled to generate beneficial effects and limit pathological differentiation. Much research over the past 20 years has identified many different molecular species that are relevant, but these have mainly been studied one at a time. In this study, we developed and implemented a novel computational strategy to interrogate the key players in this transformation process to identify which are the major bottlenecks. We determined that NFATc1 and pSP1 are essential for promoting epithelial or mesenchymal differentiation, respectively. We then predicted the existence of a partially transformed cell that exhibits both epithelial and mesenchymal characteristics. We found this partial cell type develops a network of invasive but stunted vascular structures that may be a unique cell target for understanding cancer progression and angiogenesis.

## Introduction

The epithelial to mesenchymal transition (EMT) is a broadly participating, evolutionarily conserved differentiation program essential for tissue morphogenesis, remodeling and pathological processes such as cancer (Thiery, 2003). During EMT polarized, tightly adhered epithelial cell monolayers are transformed into non-interacting motile mesenchymal cells that simultaneously degrade and synthesize extracellular matrix (ECM) components and invade into the underlying tissue space (Stahl & Felsen, 2001). EMT is the fundamental initiator of developmental processes such as embryonic gastrulation and valvulo-genesis (Eisenberg & Markwald, 1995) (also Kalluri J Clin Invest 2009, Thiery Cell 2009). Transforming growth factor *β* (TGF-*β*) family members are important inducers of both developmental and pathological EMT (Xu *et al.*, 2009, Zavadil & Böttinger, 2005). Decades of research has focused on identifying molecular regulators of EMT, but almost all on a single gene and in a nearly binary yes/no level of qualitative understanding. Medici and coworkers identified a core signaling program by which TGF-*β* isoforms induce EMT across a variety of cell lines (Medici *et al.*, 2006, 2008). This program involves carefully orchestrated rounds of gene expression driven by the Smad and Snail families of transcription factors as well as other key factors such as lymphoid enhancer-binding factor 1 (LEF-1), nuclear factor of activated T-cells, cytoplasmic 1 (NFATc1), and specificity protein 1 (Sp1). Coregulators such as *β*-catenin, NF-_κ_B, and the ErbB family of receptor tyrosine kinases however also participate in EMT regulation, but the degree of each’s influence is difficult to ascertain in isolation (Hardy *et al.*, 2010, Huber *et al.*, 2004, Jiang *et al.*, 2007, Kim *et al.*, 2002). EMT also exhibits complex temporal dynamics that are often intractable in gain/loss of function studies. Elucidating the master regulatory architecture controlling EMT therefore requires inclusion of these complex overlapping and non-binary behaviors.

Systems biology and mathematical modeling are essential tools for understanding complex developmental programs like EMT (Ahmed & Nawshad, 2007). Previous computational models of TGF–*β* induced differentiation focused on single biological factors or EMT in single cells. For example, Chung *et al.*, constructed a model of TGF-*β* receptor activation and Smad signaling using ordinary differential equations and mass-action kinetics. Their model suggested that a reduction of functional TGF-*β* receptors in cancer cells may lead to an attenuated Smad2 signal (Chung *et al.*, 2009). Similarly, Vilar *et al.* suggested that specific changes in receptor trafficking patterns could lead to phenotypes that favor tumorigenesis (Vilar *et al.*, 2006). Although these models provided insight into the role of receptor dynamics, EMT induction involves many other components, including competing second messengers and interconnected transcriptional regulatory loops. Integrating these additional scales of molecular signaling while maintaining the capacity for robust prediction requires a new and expanded computational and experimental strategy. Data-driven systems approaches (Cirit & Haugh, 2012) or logical model formulations (Morris *et al.*, 2011) are emerging paradigms that constrain model complexity through the incorporation of training and validation data. These are interesting techniques because the data informs model structure (which can be expanded as more data becomes available). Alternatively, Bailey proposed more than a decade ago that a qualitative understanding of a complex biological system should not require complete definition of its structural and parametric content (Bailey, 2001). Shortly thereafter, Sethna and coworkers showed that complex model behavior is often controlled by only a few parameter combinations, a characteristic seemingly universal to multi-parameter models referred to as “sloppiness” (Machta *et al.*, 2013). Thus, reasonable model predictions are often possible with only limited parameter information. Taking advantage of this property, we developed sloppy techniques for parameter identification using ensembles of deterministic models (Song *et al.*, 2010). Furthermore, we proposed that the sloppy behavior of biological networks may also be seen as a source of cell-to-cell (Lequieu *et al.*, 2011) or even patient-to-patient heterogeneity (Luan *et al.*, 2010). Bayesian parameter identification techniques have also been used to explore cell-to-cell heterogeneity (Hasenauer *et al.*, 2011, Kalita *et al.*, 2011), where a population of cells could be viewed as a dynamic ensemble of context-specific biochemical networks (Creixell *et al.*, 2012).

In this study, we developed a family of mathematical models describing the induction of EMT by TGF–*β* isoforms in the presence and absence of vascular endothelial growth factor A (VEGF-A). We integrated a simple rule-based description of activity and gene expression regulation with traditional ordinary differential equation (ODE) modeling to describe an EMT interaction network containing 97 gene, protein or mRNA components interconnected through 169 interactions. This integration allows the description of complex regulatory interactions in the absence of specific mechanistic information, it also allowed to build a predictive yet compact model. A family of model parameters was estimated using 41 molecular data sets generated in DLD1 colon carcinoma, MDCKII and A375 melanoma cells using the Pareto optimal ensemble technique (JuPOETs) multiobjective optimization algorithm. JuPOETs generated an ensemble of approximately 1400 models for analysis. Analysis of the model population suggested that both MCF10A and DLD1 cells could exhibit phenotypic heterogeneity if treated simultaneously with TGF-*β*1/2 and VEGF-A. This heterogeneity was characterized by a significant fraction of the population being in a “hybrid state” having both high E-cadherin and high Vimentin expression. We tested these predictions using qRT-PCR and flow-cytometry studies in a variety of experimental conditions. Validation studies confirmed that upwards of 45% of the cellular population could be put into the hybrid state in the presence of both TGF-*β*1/2 and VEGF-A. Moreover, this response depended upon both activation of Sp1 by MAPK and NFATc1 transcriptional activity consistent with the predicted molecular signaling. Lastly, the hybrid populations of both DLD1 and MCF10A cells exhibited different functional behavior than those from either TGF-*β* or VEGF-A treatment. The extent of ductal branch formation significantly increased with MCF10A cells in the hybrid phenotype, compared with cells treated with VEGF-A alone. Together, these results establish a predictive mechanistic model of EMT susceptibility, and reveal a novel signaling axis, which possibly regulates carcinoma progression through an EMT versus tubulogenesis response.

## Results

### The model population captured key features of TGF-*β* induced EMT

The EMT model architecture, based upon curated molecular connectivity, described the expression of 23 genes following exposure to TGF-*β* isoforms and VEGF-A (Fig. 1). The EMT model contained 74 molecular species interconnected by 169 interactions. Model equations were formulated as ordinary differential equations (ODEs) augmented with rule-based descriptions of activity and gene expression regulation. ODEs are common tools to model biochemical pathways (Chen *et al.*, 2009, Schoeberl *et al.*, 2002, Tasseff *et al.*, 2011). However, while ODE models can simulate complex intracellular behavior, they require estimates for model parameters which are often difficult to obtain. The EMT model had 251 unknown model parameters, 169 kinetic constants 38 control constants and 44 saturation constants. As expected, these parameters were not uniquely identifiable given the training data (Gadkar *et al.*, 2005). Thus, instead of identifying a single best fit (but uncertain) model, we estimated a sloppy population of models (each consistent with the training data) by simultaneously minimizing the difference between model simulations and 41 molecular data sets using the Pareto Optimal Ensemble Technique (JuPOETs). The training data were generated in DLD1 colon carcinoma, MDCKII, and A375 melanoma cells following exposure to TGF-*β* isoforms (Medici *et al.*, 2008). We organized these data sets into 11 objective functions which were simultaneously minimized by JuPOETs. Additionally, we used data generated in this study (Fig. S4), and 12 molecular data sets generated in HK-2 cells following VEGF-A exposure to train VEGF-A responsive model processes (Lian *et al.*, 2011). To guard against overfitting, we augmented the multiobjective optimization with leave-one-out cross validation to independently estimate both the training and prediction error for each objective. Thus, we generated 11 different model ensembles. Lastly, we compared model predictions with independent data sets not used during training (both at the molecular and model population levels) to evaluate the predictive power of the parameter ensemble.

**Fig. 1:**
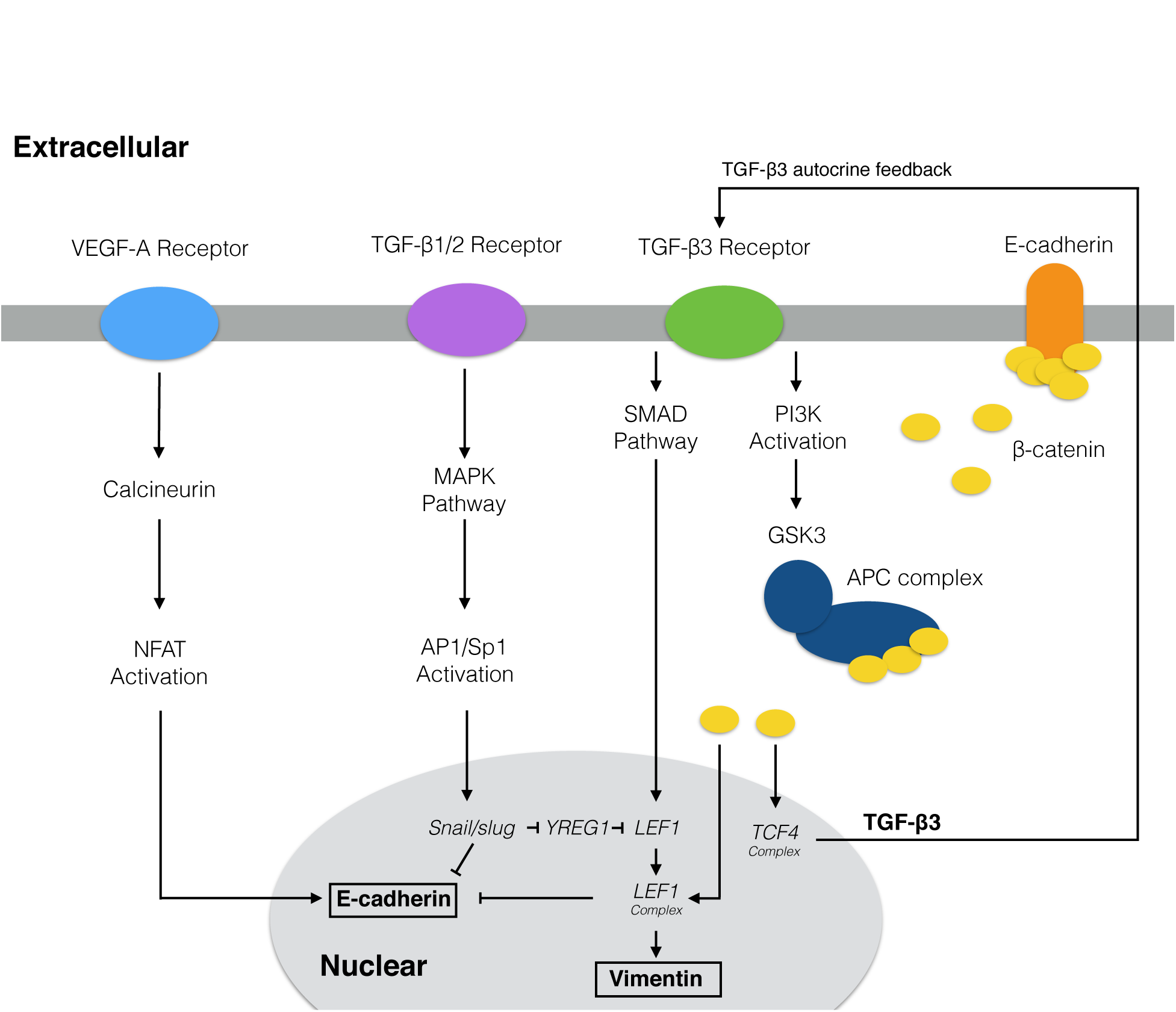
Model connectivity recreates the core architecture during EMT. The EMT network contains 97 nodes (proteins, mRNA, and genes) interconnected by 169 interactions. Central to EMT induction, activation of the MAPK cascade occurs through TGF-*β*1/2 binding which activates the AP-1/Sp1 transcriptional axis. AP-1/Sp1 drives an autocrine response of TGF-*β*3, which activates the Smad cascade, leading to phenotypic change. Conversely, VEGF-A binding can stabalize an epithelial phenotype through NFAT activation. Downstream activation of /3-catenin signaling due to E-cadherin loss and GSK3 inactivation of *β*-cateinin confinement is critical to the complete activation of the EMT program. The complete list of molecular interactions that comprise the model is given in the supplement.

JuPOETs generated a population of probable signaling models which captured the multiple phases of EMT induction (Fig. 2). JuPOETs sampled well over 10^4^ probable models during each stage of the cross-validation using global random sampling. From this analysis, N ≃ 1400 models were selected for further analysis. The selected models all had the same possible molecular connectivity, but different values for model parameters. Transcription and translation rates, as well as mRNA and protein degradation terms, were set using physical values from the literature (Milo *et al.*, 2010), and allowed to vary by a scaling factor, see methods. Model selection was based upon Pareto rank, the prediction and training error across all objectives. The model population recapitulated key signaling events following TGF-*β* exposure. We subdivided the response to TGF-*β* exposure into two phases. First, TGF-*β*1/2 signaling initiated a program which down-regulated E-cadeherin expression in a MAPK dependent manner while simultaneously upregulating TGF-*β*3 expression. Second, TGF-*β*3 secretion initiated an autocrine feedback which upregulated the expression of mesenchymal markers such as Vimentin and key upstream transcription factors such as LEF-1 in a SMAD dependent manner. TGF-*β*3 expression was also able to sustain *β*-catenin release by inhibiting its sequestration by the APC complex through PI3K mediated GSK3, which was captured by the model (Fig. 4B). Each phase involved the hierarchal expression and/or post-translational modification of several key transcription factors. During the first phase, stimulation with TGF-*β*1/2 (10 a.u.) activated both the SMAD and MAPK pathways. MAPK activation resulted in the phosphorylation of the transcription factor activator protein 1 (AP-1), which in-turn upregulated the expression of Snail, a well established transcriptional repressor (Fig. 2A). Snail expression was MAPK-dependent; the MEK inhibitor U0126 blocked AP-1 activation and Snail expression following TGF-*β*1/2 exposure (Fig. 2A, Lane 3). Similar results were obtained for Slug expression, confirming initial activation through the MAPK pathway (data not shown). Overexpression of either Snail or Slug upregulated TGF-*β*3 expression (Fig. 2C) while simultaneously downregulating E-cadeherin expression (Fig. 2F). During the second phase, TGF-*β*3 secretion and the subsequent autocrine signaling resulted in the upregulation of mesenchymal marker expression. The TGF-*β*3 induced gene expression program involves a complex hierarchy of transcriptional and post-translational regulatory events. Absence of E-cadherin indirectly promoted TGF-*β*3 expression through the *β*-catenin/TCF4 complex following Snail or Slug expression (Fig. 2C, Lane 2 or 3). Conversely, over-expression of E-cadherin inhibited the TGF-*β*3 autocrine production by sequestering cytosolic *β*-catenin, thereby blocking EMT (Fig. 2C, Lane 4 or 5). TGF-*β*3 signaled through the Smad pathway to regulate LEF-1 expression and downstream target EMT genes (Fig. 2G). TGF-*β*3 (10 a.u.) in combination with downstream inhibitors (DN-Smad4 and DN-LEF-1) completely inhibited Vimentin expression, while elevating E-cadherin expression (Fig. 2H,I).

**Fig. 2:**
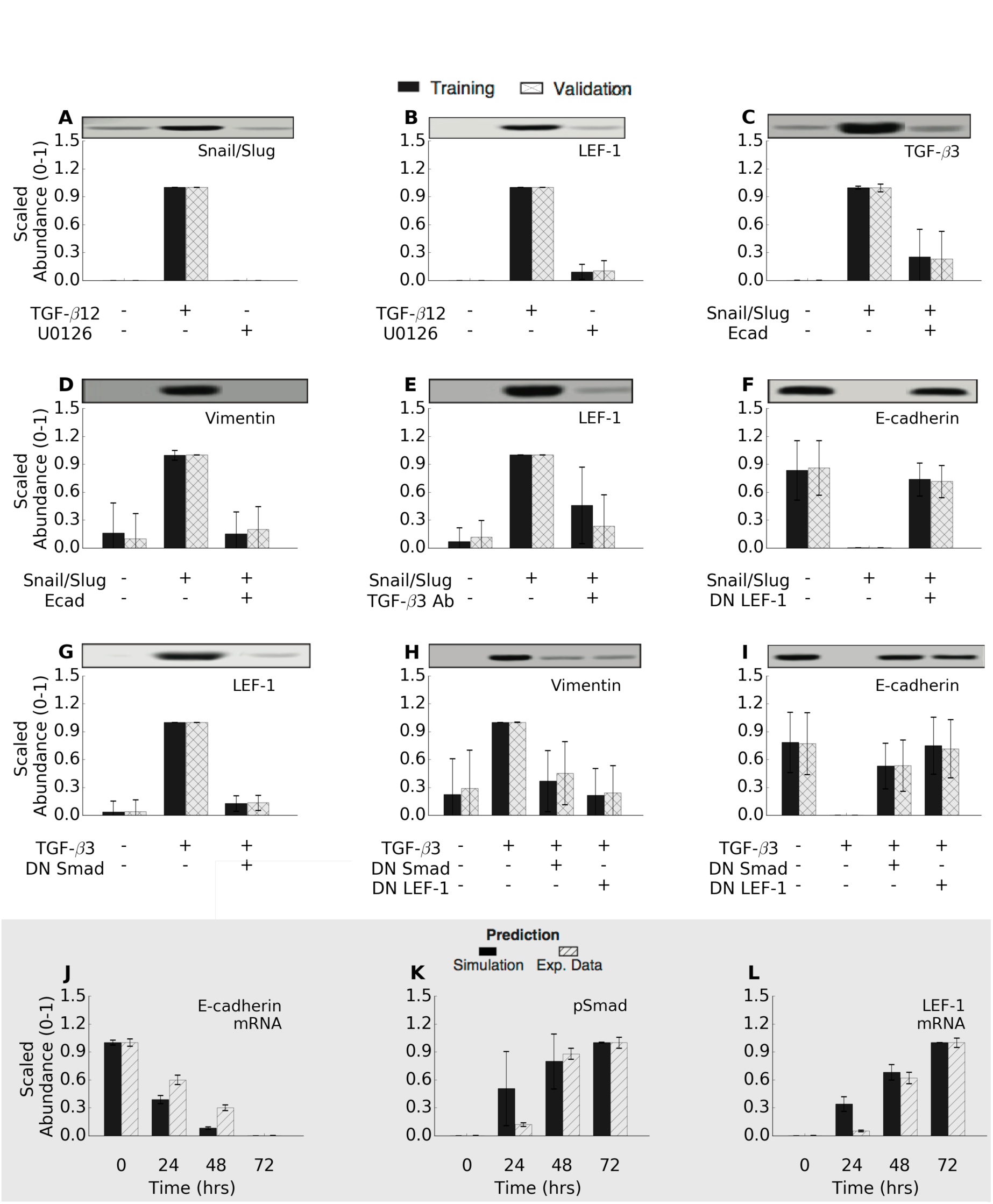
Training and validation simulations. The population of EMT models qualitatively captured TGF-*β*-induced EMT signaling. (A-I) The population was generated using JuPOETs and trained using 11 different objective functions (41 data sets) taken from Medici *et al.* (Medici *et al.*, 2008). The model captured the simulated experiments for all cases when compared to randomized controls. (J-L) The model populations were also compared against untrained temporal data to measure the effectiveness as a pure prediction.

The predictive power of the ensemble was tested using cross validation and by comparing simulations with data not used for model training. In whole, all of our training objectives were statistically significant (at a 95% confidence interval) compared to a randomized parameter family (N = 100) generated from a random starting point. Conversely, we *predicted* all of the training objectives, at a 95% confidence interval compared to randomized parameters (Wicoxon non-parametric test). The model also captured the temporal gene expression responses of E-cadherin, pSmad2, and LEF-1 (not used for model training) to within one-standard deviation (up to the 48 hr time-point) (Fig. 2J-L). Taken together, the model captured the key signaling events revealed by Medici *et al.* (Medici *et al.*, 2008) that drive the phenotypic conversion. A listing of objective function values resulting from training, cross validation and the random parameter control is given in the supplement (Fig. S1).

### Identification of a novel LEF-1 regulator

During model identification, we found that consistent TGF-*β* induced EMT from a stable epithelial cell population required an additional regulatory protein. This protein, which we called hypothetical regulator 1 (YREG1), was required to mediate between SNAIL/SLUG transcriptional activity and the upregulation of LEF-1 expression following TGF-*β*1/2 exposure. SNAIL/SLUG are well known transcriptional repressors (Dhasarathy *et al.*, 2011, Hemavathy *et al.*, 2000a,b), although there are a few studies which suggest that at least SNAIL can also act as a transcriptional activator (Guaita *et al.*, 2002). In the model, we assumed the expression of SNAIL/SLUG was likely regulated by AP1/SP1 (Jackstadt *et al.*, 2013). Thus, upon receiving direct SNAIL/SLUG and TGF-*β*3 signals, the model predicted enhanced SNAIL/SLUG expression, consistent with experimental observations. TGF-*β*1/2 stimulation also induces LEF-1 expression. However, literature evidence suggested that LEF-1 expression was not strongly dependent upon AP1/SP1 activity (Eastman & Grosschedl, 1999). Thus, either SNAIL/SLUG are acting as inducers (contrary to substantial biochemical evidence) or, they are repressing the expression of an intermediate repressor. Given the biochemical evidence supporting SNAIL/SLUG as repressors, we created the hypothetical YREG1 repressor whose expression is downregulated by SNAIL/SLUG. The literature data therefore suggested that YREG1 had two transcriptional targets, LEF-1 and TGF-*β*3. By adding this regulator, our simulations became consistent with training and literature data. Medici *et al.* suggested that feedback between *β*-catenin and LEF-1 was likely, although this feedback had yet to be identified (Medici *et al.*, 2008). Low levels of YREG1 expression were present in all simulations to regulate the formation of the *β*-catenin-LEF-1 complex. To test the effect of YREG1 on the epithelial population, we conducted over-expression and knockdown simulations on untreated cells (Fig. 4C and 4D). In the absence of YREG1, the population of models failed to consistently to retain a stable epithelial state (Fig. 4D). Conversely, YREG1 amplification revealed an enhanced epithelial phenotype, while some inherently transformed cells moved towards a hybrid phenotype (Fig. 4C). Elevated YREG1 repressed LEF-1 and TGF-*β*3 expression, thereby not allowing free *β*-catenin to form the *β*-catenin-LEF-1 complex, or TGF-*β*3 induced SMAD activation. Taken together, low YREG1 expression was required for the maintenance of a stable epithelial phenotype that was simultaneously inducible across TGF-*β*1/2, TGF-*β*3 and SNAIl/SLUG transfection.

### TGF-*β*1/2 and VEGF-A exposure promotes phenotype heterogeneity through NFATc and phosphorylated Sp1

While we captured the central tendency of many of the molecular features of EMT induction following TGF-*β*1/2 exposure, an often neglected but important emergent feature of developmental and pathological programs is population heterogeneity (Park *et al.*, 2010). We (and others) have previously hypothesized that deterministic model ensembles can simulate population behavior, at least at a course grained level (Lequieu *et al.*, 2011). We tested this hypothesis by analyzing the response of the population of EMT models to extracellular cues and then comparing this response to flow cytometry studies. We quantified the phenotypic response of the individual members of the ensemble to TGF-*β*1/2 stimulation for two downstream phenotypic markers, Vimentin (mesenchymal) and E-cadherin (epithelial) following the addition of TGF-*β*1/2 alone (Fig. 3), and/or VEGF-A in combination with NFATc inhibitors (Fig. 3).

**Fig. 3:**
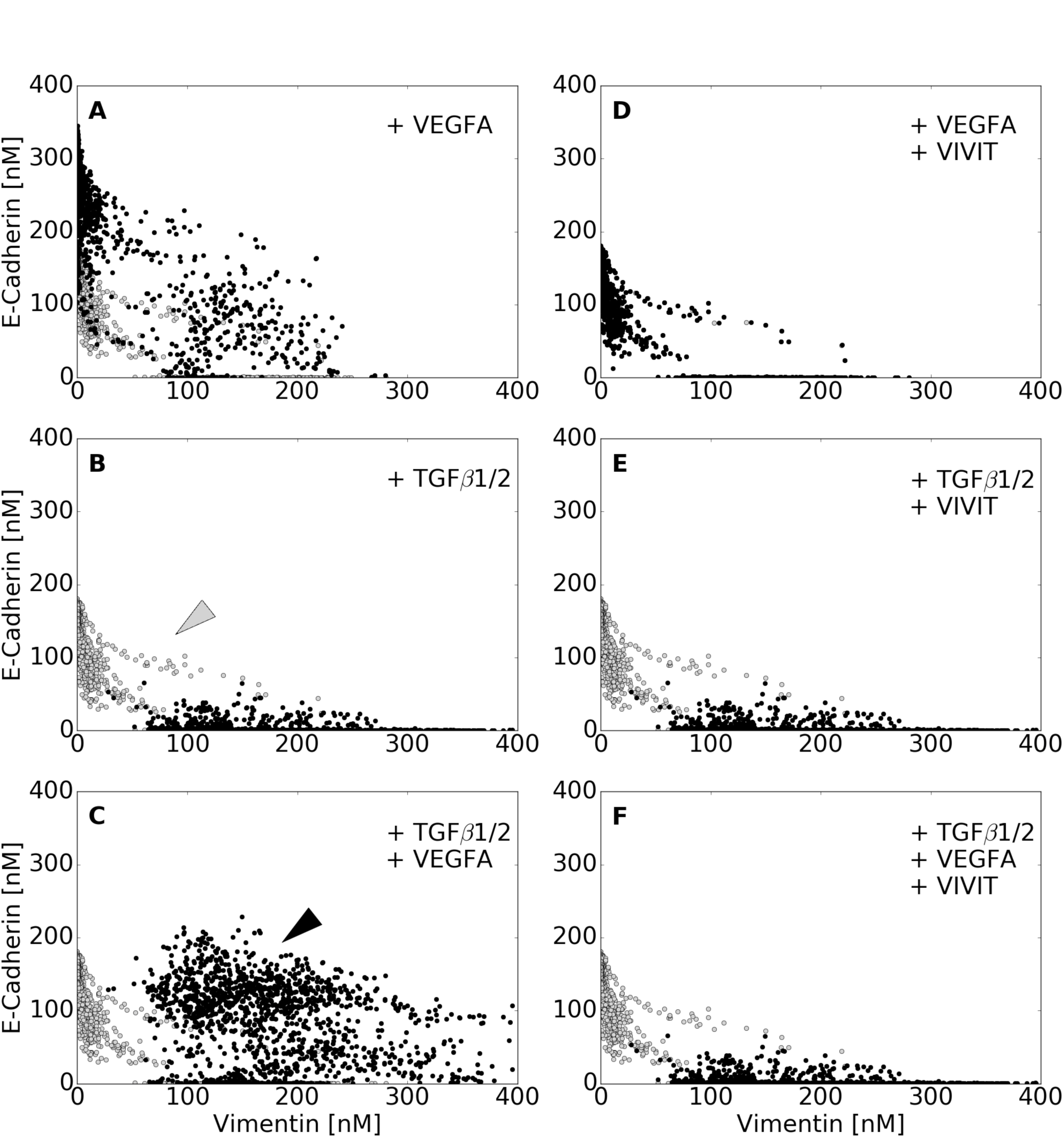
Simulated VEGF-A and TGF-*β*1/2 exposure promoted phenotype heterogeneity. Simulated response to TGF-*β*1/2 and VEGF-A exposure with and without axis specific inhibitors. Vimentin and E-cadherin abundances (in nM) were used to quantify the shift in population at 48 hrs. (A-C) VEGF-A (50 a.u.) treatment resulted in a population with enhanced epithelial properties. This was contrary to the addition of TGF-*β*2 (10 a.u.), which shifted the population towards a mesenchymal phenotype. Interestingly, the combined effects of TGF-*β*2 and VEGFA was found to increase both ecadherin and vimentin levels, creating a heterogeneous population (black arrow), which can also be seen in a minority of untreated cells (gray arrow). (D-F) To isolate the effect of NFAT, we inhibited NFAT de-phosphorylation in combination with VEGFA. This negated the increase in ecadherin expression and shifted the population towards a mesenchymal phenotype. Likewise, combining NFAT inhibition with TGF-*β* mitigated all VEGF enhanced ecadherin expression. Lastly, combination of TGF-*β*2, VEGFA, and NFAT inhibition nearly mitigated all effects of VEGFA, shifting the heterogeneous population towards a mesenchymal phenotype. In whole, high levels of phosphorylated-Sp1 correlated with vimentin expression, while NFAT was responsible for maintaining E-cadherin expression in the presence of other factors, although neither were mutually exclusive.

We identified model subpopulations that exhibited different behaviors following exposure to TGF-*β*1/2 (Fig. 3B). Analysis of the molecular signatures of these subpopulations suggested the abundance, localization and state of the Sp1, AP-1 and NFATc transcription factors controlled population heterogeneity. The majority of models (>80%) responded to treatment, moving away from the untreated population (Fig 3A-F, gray). These models showed the classically expected behavior, a switch from an epithelial to mesenchymal phenotype following TGF-*β*1/2 exposure. Some models resembled untreated cells; they had elevated phosphorylated Sp1, relative to non-induced cells, which decreased E-cadherin expression through Slug-mediated inhibition, which in turn increased Vimentin expression through TGF-*β*3 autocrine signaling and the liberation of *β*-catenin. However, the most biologically interesting behavior was exhibited by cells achieving a hybrid phenotype, most notable in a dual treatment condition (3C, black arrow), but also present in a small percentage of untreated cells (Fig. 3B, gray arrow). Models with this hybrid phenotype had elevated Sp1 and NFAT transcriptional activity, resulting in simultaneously increased Vimentin and E-cadherin expression (Fig. 4A).

**Fig. 4:**
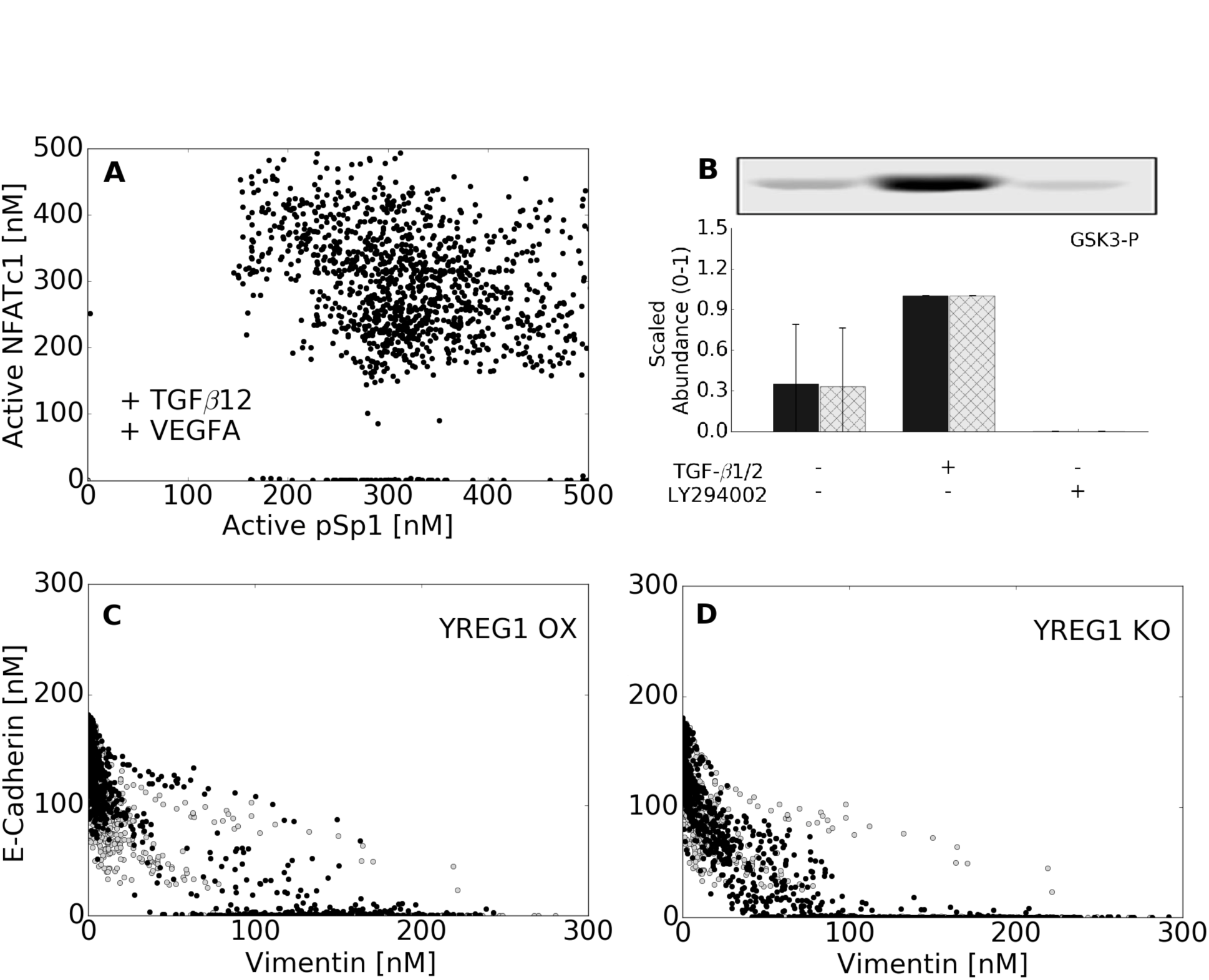
Analysis of underlying signaling responses. (A) We examined the distribution of NFATc1 and AP1/SP1 in cells containing the hybrid phenotype (VEGF-A + TGF-*β*2 case), showing the potential for cells to express both SP1 and NFATc1 in a non exclusive manner. (B) We were able to show a fit to an additional objective demonstrating the actiation of GSK3 through PI3K. Our model captures this activation through TGF-*β*3 signaling. LY294002 is a PI3K inhibitor. (C) We identified a novel regulator of LEF1 called YREG1 that allows Snail/Slug to emulate an inducer by repressing YREG1, which was required to stabalize the untreated population. YREG1 overexpression revealed an enhanced epithelial phenotype, while some inherently transformed cells moved towards a hybrid phenotype. (D) In the absence of YREG1, most of the population failed to consistently to retain a stable epithelial state.

To better understand the hybrid phenotype, we simulated the response of the model population to TGF-*β*1/2 and VEGF-A treatment with and without NFATc inhibitors (Fig. 3). As expected, stimulation with VEGF-A (50 a.u.) maintained an epithelial population (Fig. 3A), while TGF-*β*1/2 (10 a.u.) exposure shifted the population from an epithelial to a mesenchymal phenotype (Fig. 3B). On the other hand, combined stimulation with TGF-*β*1/2 (10 a.u.) and VEGF-A (50 a.u.) increased both E-cadherin and Vimentin expression, resulting in a hybrid phenotype with both epithelial and mesenchymal characteristics (Fig. 3C). Vimentin expression was correlated with high levels of nuclear phosphorylated Sp1, following TGF-*β*1/2 exposure. Conversely, elevated E-cadherin expression depended upon the activity of NFAT transcription factors downstream of VEGF-A stimulation. To further isolate the role of NFAT on this hybrid state, we simulated the inhibition of NFAT transcriptional activity across all conditions (all else being equal). NFAT inhibition in combination with VEGF-A or TGF-*β*1/2 treatments blocked increased E-cadherin expression in the case of VEGF-A (Fig. 3D), but did not influence TGF-*β*1/2 signaling (Fig. 3E). Lastly, NFATc inhibition in combination with simultaneous TGF-*β*1/2 and VEGF-A exposure repressed nearly all E-cadherin expression, shifting nearly the entire population towards a mesenchymal phenotype (Fig. 3F). Taken together, high levels of nuclear localized phosphorylated Sp1 correlated with Vimentin expression, while NFATc transcriptional activity was critical for maintaining E-cadherin expression in the presence of competing signals.

### Combined TGF-*β*2 and VEGF-A exposure drives heterogeneity in MCF10A and DLD1 cells

The EMT model simulations suggested the transcriptional activity of NFATc and Sp1 could be independently tuned to generate a hybrid cell population with both epithelial and mesenchymal characteristics. To test this hypothesis, we exposed either quiescent epithelial (MCFA10, Fig. 5) or transformed epithelial cells (DLD1, Fig. S2) to combinations of TGF-*β*1/2 and/or VEGF-A. As expected, TGF-*β*1/2 treatment (10ng/ml) increased Slug and Vimentin expression, while repressing E-cadherin expression both at the transcript and protein levels in MCF10A (Fig. 5A-B) and DLD1 cells (Fig. S3C). Both MCF10A (Fig. 5C) and DLD1 cells (Fig. S2E,G) transitioned from quiescent cobblestone morphology to spread spindle shapes, consistent with EMT. As predicted, we found increased nuclear localization of phosphorylated Sp1 following TGF-*β*1/2 stimulation in both MCF10A (Fig. 5B,C) and DLD1 cells (Fig. S2E,F). Consistent with model predictions, VEGF-A (50ng/ml) treatment increased the abundance of NFATc1 and E-cadherin at both the transcript and protein level in both MCF10A (Fig. 5A) and DLD1 cells (Fig. S2). We also found that NFATc1 nuclear localization significantly increased in both MCF10 (Fig. 5B,C) and DLD1 (Fig. S2C,E) cells treated with VEGF-A. Interestingly, combining VEGF-A (50ng/ml) with TGF-*β*1/2 (10ng/ml) resulted in significantly elevated expression of both E-cadherin and Vimentin at the transcript and protein levels in both MCF10A (Fig. 5A,B) and DLD1 cells (Fig. S2–S3). NFATc1 expression increased, while Sp1 expression was similar to the TGF-*β*1/2 case alone (Fig. 5A-B and Fig. S2D,E), supporting their independent regulation. The expression of Slug, and Vimentin significantly increased, while E-cadherin levels were increased in MCF10A cells (Fig. 5A) and maintained at control levels in DLD1 cells (Fig. S2D). As predicted, nuclear co-localization of both NFATc1 and phosphorylated Sp1 were apparent in MCF10A (Fig. 5B,C) and DLD1 (Fig. S2E,F) cells treated with both ligands. Taken together, combined VEGF-A and TGF-*β*1/2 treatment elicited a hybrid phenotype expressing both mesenchymal and epithelial characteristics in both MCF10A and DLD1 cells. This phenotype was driven by the transcriptional activity of two key transcription factors, Sp1 and NFATc, which could be modulated independently by TGF-*β*1/2 and VEGF-A exposure.

**Fig. 5:**
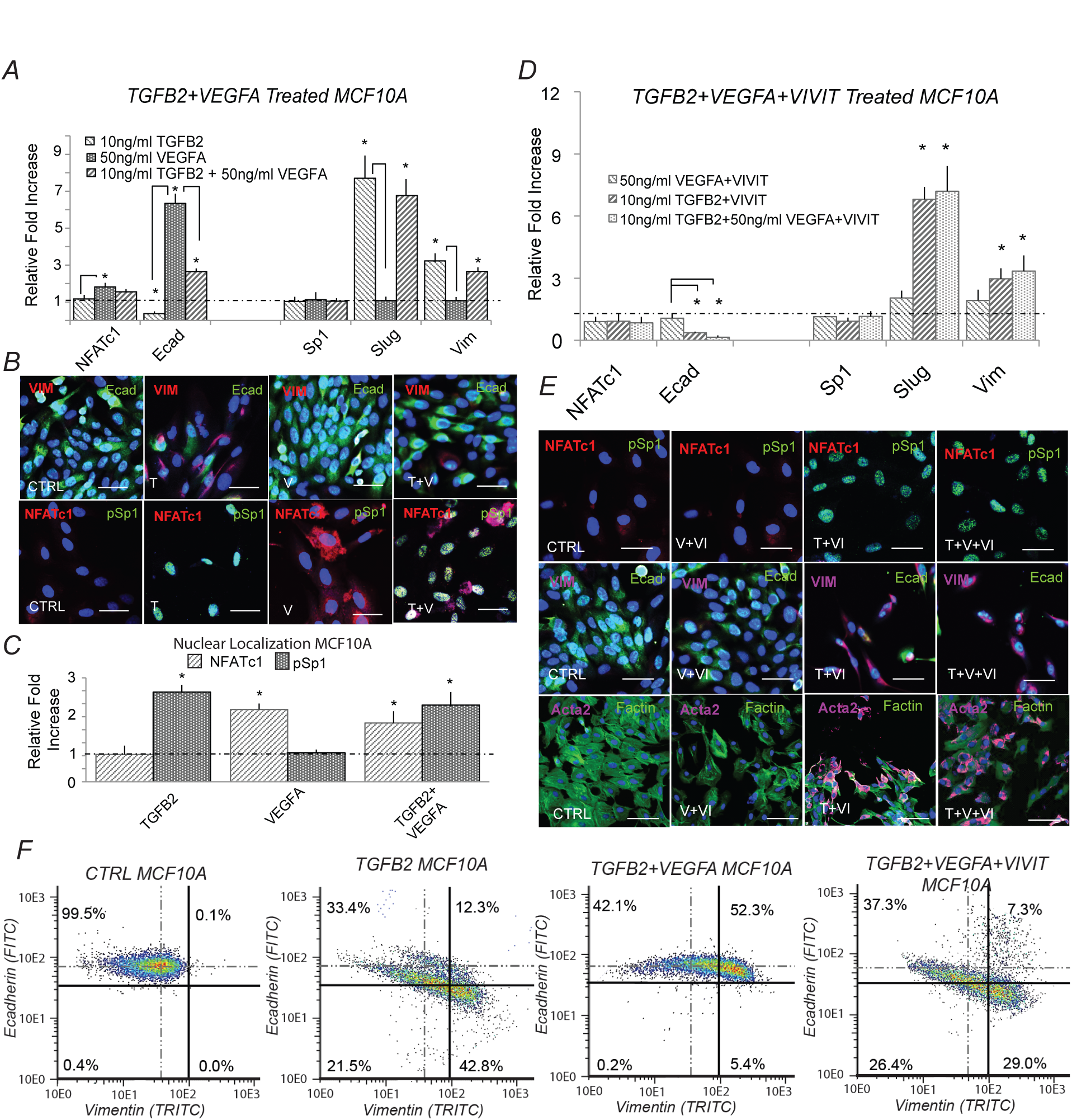
Simultaneous TGF-*β*1/2 and VEGF-A treatment induced phenotype heterogeneity and is dependent upon NFAT activity in-vitro. (A) In MCF10A, treatment with (10ng/ml) TGF-*β*2 increased Slug and vimentin, while ecadherin expression was inhibited at both the gene and protein level at 48 hrs. Conversely, VEGFA alone increased both NFATc1 and ecadherin gene expression. Simultaneous TGF-*β*2 (10ng/ml) and VEGFA (50ng/ml) treatment increased Slug, NFATc1, and vimentin expression, while also increasing ecadherin levels via qPCR. (B-C) Immunofluorescence confirmed these results and nuclear co-localization of both phospho-Sp1 and NFAT were found dependent upon TGF-*β*2 and VEGFA, respectively. (D) To isolate the effect of NFAT, treatment of VEGFA (50ng/ml) and VIVIT (10*μ*M) reduced ecadherin expression at 48hrs (control-dashed line). Similarly, combined TGF-*β*2, VEGFA and VIVIT treatment increased Slug and vimentin expression, while inhibiting ecadherin levels via qPCR. (E) These findings were confirmed via immunofluorescence as the VIVIT peptide inhibited ecadherin and nuclear localization of NFATc1 in all three cases. (F) Quantitative flow cytometry also confirmed this trend. Similar experiments in DLD1 followed a similar trend (supplement). Magnification, 40x. Scale bars: 50*μ*m. C=Control, T=TGF-*β*2, V=VEGFA, VI= NFAT inhibitor (VIVIT). Asterisks signify statistical differences from each other according to a one-way ANOVA with Tukey’s post hoc (p≺0.05).

Our phenotypic analysis predicted that NFATc transcriptional activity was critical to maintaining E-cadherin expression in the presence of both VEGF-A and TGF-*β*1/2. We experimentally tested this hypothesis by exposing both MCF10A (Fig. 5E,F) and DLD1 cells (Fig. S3) to combinations of VEGF-A and TGF-*β*1/2 in the presence or absence of VIVIT, a soluble peptide inhibitor of NFATc transcriptional activity (Aramburu *et al.*, 1999). Treatment with VEGF-A (50ng/ml) and VIVIT (10*μ*M) in MCF10A cells significantly reduced E-cadherin expression compared to VEGF-A alone (Fig. 5D,E). Co-treatment with VIVIT and TGF-*β*1/2 did not enhance EMT capacity of MCF10A cells above that of TGF-*β*1/2 alone (Fig. 5A,B,E). Likewise, VIVIT in combination with both TGF-*β*1/2 and VEGF-A resulted in a loss of E-cadherin gene and protein expression, while Slug and Vimentin levels remained increased (Fig. 5D,E). Quantitative flow cytometry confirmed these results in both MCF10A (Fig. 5F) and DLD1 cells (Fig. S3C). Both epithelial cell lines initially had high levels of E-cadherin expression, and low Vimentin abundance (Q1-99.5%), but both MCF10A and DLD1 cells shifted from an epithelial to mesenchymal phenotype (Q1-33.4%, Q4-42.8%) following TGF-*β*1/2 exposure. As expected, NFATc nuclear localization was repressed with VIVIT treatment regardless of ligand stimulation, while the abundance of nuclear phosphorylated Sp1 increased for both TGF-*β*1/2 and TGF-*β*1/2 + VIVIT conditions (Fig. 5C,E). Combined TGF-*β*1/2 and VEGF-A increased both Vimentin and E-cadherin expression (Q1-42.1%, Q2-52.3%) compared to TGF-*β*1/2 alone. Together, these results demonstrate that NFATc and phosphorylated Sp1 are critical for regulating E-cadherin and Vimentin expression during phenotype heterogeneity in MCF10A and DLD1.

### Ductal branching during acini formation is dependent upon phenotype heterogeneity in MCF10A and DLD1 cells

We finally employed established three-dimensional (3D) in vitro models of invasion, migration, compaction, and tubulogenesis (Dhimolea *et al.*, 2010) to determine the functional consequences of the hybrid phenotype (Fig. 6). MCF10A and DLD1 cells were aggregated via hanging drop, placed on the surface of a collagen gel, and cultured for 72 hrs under various biochemical treatments. TGF-*β*1/2 stimulation significantly enhanced cell matrix invasion and matrix compaction, while in contrast VEGF-A stimulation promoted surface migration but no invasion or compaction (Fig. 6B-D). Interestingly, combined TGF-*β*1/2 and VEGF-A stimulation significantly increased cell migration potential above that of VEGF-A alone while maintaining 3D matrix compaction, though with decreased magnitude compared to TGF-*β*1/2 alone. Inhibition of NFATc transcriptional activity by VIVIT decreased migration following treatment with VEGF-A alone (Fig. 6B). Co-treatment of VIVIT significantly decreased migration, while complementarily increasing invasion and compaction, when MCF10A cells were stimulated with both VEGF-A and TGF-*β*1/2 (Fig. 6B-D). The responses of DLD1 cells followed a similar trend to MCF10A, although the magnitudes of migration, invasion, and compaction were less. Cell circularity within 3D gels strongly and negatively correlated with both invasion and compaction regardless of treatment (Fig. 6E). Circularity refers to the morphology of the cells. In general, a quiescent epithelial cells assumes a circular morphology in culture, while an active mesenchymal cell is highly elongated. The circularity index, a common means of quantifying cell morphology, relates cell area to perimeter. A perfect circle has a circularity index equal to 1.0, while a straight line has a circularity index equal to 0.0, see Butcher *et al.* (Butcher *et al.*, 2004). TGF-*β*1/2 treatment alone resulted in irregular and spindle shaped morphology, while VEGF-A exposure promoted round quiescent cells (Fig. 6A). Combined VEGF-A and TGF-*β*1/2 promoted morphology between these extremes. VIVIT mediated NFATc inhibition significantly reduced the circularity index, similar to TGF-*β*1/2 treatment (Fig. 6F). VEGF-A treatment also induced the formation of tubular structures (acini), but the number of tubular branches relative to total acini was significantly increased upon combined TGF-*β*1/2 and VEGF-A. No tubular structures were identified within the DLD1 constructs during the 7 day tubulogenesis end-points, supporting that MCF10A and DLD1 cells have some cell-type specific EMT sensitivity despite their underlying competency for acquiring a heterogeneous phenotype. This suggests that initial EMT sensitivity of a cell influences downstream functional response from TGF-b and VEGFA stimulation. Together, these results establish that VEGF-A and TGF-*β*1/2 ligand concentrations potentiate between acini and ductal branch formation in 3D culture, and are dependent upon NFATc activity.

**Fig. 6:**
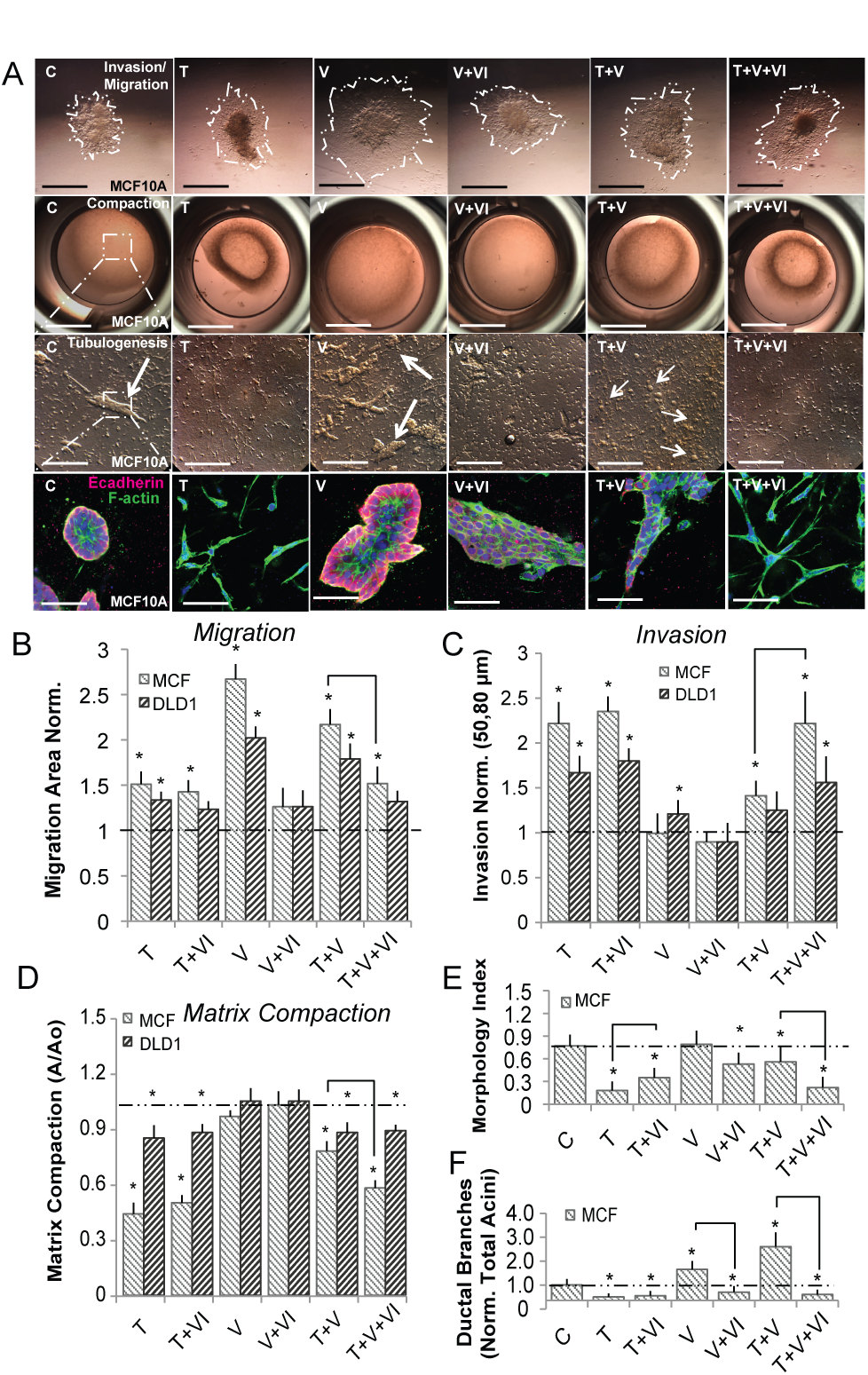
Ductal branching is dependent upon phenotype heterogeneity within MCF10A in 3-D culture. MCF10A and DLD1 were formed into spheroids overnight and explanted to a collagen gel for 72 hrs. For compaction and tubular assays, cells were embedded into collagen gels for 72 hrs, and the extent of tubulogenesis was measured at 7 days. (A-D) Within MCF10A, TGF-*β*2 (10ng/ml) enhanced invasion and contractile properties while, VEGFA (50ng/ml) promoted increased migration. TGF-*β*2 with VEGFA significantly increased migration, while limiting with compaction. VIVIT (10*μ*M) in combination with VEGFA and TGF-*β*2 decreased migration and compaction, while increasing invasion. (D) Likewise, cell morphology (circularity index) correlated with both invasion and compaction in MCF10A. (E-F) The size of tubular structures (acini) also increased significantly upon addition of VEGFA, while the number of ductal branches was most significant upon simultaneous TGF-*β*2 and VEGFA treatment (Red-Ecadherin, Green-Factin, Blue-Nuclear). DLD1 cells followed a similar trend, although the degree of migration, invasion, and compaction was less significant. In addition, no tubular structures were identified during the 7 day tubulogenesis end-points. Scale bars: 500*μ*m, 1000*μ*m, 250*μ*m, and 80*μ*m, respectively. C=Control, T=TGF-*β*2, V=VEGFA, VI= NFAT inhibitor (VIVIT). Asterisks signify statistical differences from each other according to a one-way ANOVA with Tukey’s post hoc (p≺0.05). Boxes in the left-most panel identify regions identified by arrows that were then imaged in greater zoom in the panel immediately below. The box diagram was not repeated for arrows in the other panels for clarity, but the same method was applied.

## Discussion

In this study, we developed a family of mathematical models describing the induction of EMT by TGF-*β* isoforms in the presence and absence of VEGF-A. The model, which contained 74 molecular species interconnected by 169 interactions, described the expression of 23 genes in response to growth factor stimulation. We estimated an ensemble of likely model parameters using the JuPOETs multiobjective optimization framework. The model population was trained and cross-validated to prescribe biological significance using 41 data sets generated in DLD1 colon carcinoma, MDCKII, and A375 melanoma cell lines (Medici *et al.*, 2008). Analysis of this population predicted possible phenotypic modes (and their associated signaling) that cells could exhibit when stimulated with TGF-*β* and/or VEGF-A. The most novel hypothesis generated from the analysis was that cells could operate in a hybrid state defined by both epithelial and mesenchymal traits when stimulated simultaneously with TGF-*β* and VEGF-A. We tested this hypothesis in MCF10A and DLD1 cells stimulated with combinations of TGF-*β* and VEGF-A. As expected, in the presence of TGF-*β* or VEGF-A alone, MCF10A and DLD1 cells were either mesenchymal or epithelial, respectively. However, with both TGF-*β* and VEGF-A, MCF10A and DLD1 cells exhibited a hybrid phenotype, having both epithelial and mesenchymal characteristics. Furthermore, we found that functional traits such as tubulogenesis and ductal branching were different for cells in this hybrid phenotype. Together, this study established a predictive model of EMT induction, determined that deterministic model ensembles could predict population heterogeneity, and proved the existence of a unique hybrid phenotype resulting from the simultaneous integration of extracellular growth factor signals.

Cells routinely process a multitude of signals simultaneously, especially when coordinating developmental or pathological programs. For example, oncogenic cells integrate both mechanical and chemical cues in their local microenvironment during tumorigenesis, including cytokines VEGF and TGF-*β* (Hong *et al.*, 2013). VEGF-A mediates pathological angiogenic remodeling of tumors (Nagy *et al.*, 2007), while TGF-*β* can elicit both protective and oncogenic responses (Ferrara, 2002, Willis & Borok, 2007). While much research has tested signaling pathways individually, far less is understood about combinatorial stimulation, such as with both VEGF-A and TGF-*β.* Both in vitro and in vivo studies have suggested that epithelial cells can exhibit heterogeneous phenotypes in addition to classically defined epithelial or mesenchymal states (Polyak & Weinberg, 2009, Strauss *et al.*, 2011). For example, expression profiling in human epithelial cancer cell lines demonstrated a spectrum of phenotypes, including some that expressed both E-cadherin and Vimentin simultaneously (Neve *et al.*, 2006, Welch-Reardon *et al.*, 2014). Zajchowski *et al.*, speculated that these expression profiles were somehow important for maintaining epithelial properties, while simultaneously allowing other functional behavior such as proliferation and migration (Zajchowski *et al.*, 2001). Whether and how heterogeneous phenotypes arise and participate in cancer progression, as well as their response to pharmacological inhibition are fundamental questions that should receive increased attention. In this study, we determined that a hybrid phenotype could be obtained through combined treatment with VEGF-A and TGF-*β*, both common factors localized in the tumor microenvironment. Furthermore, our systematic simulation-experimentation strategy identified that the transcriptional activity of Sp1 and NFATc were the critical factors controlling this phenotypic heterogeneity. Several studies have highlighted the importance of NFATc as a key transcription factor involved in cell growth, survival, invasion, angiogenesis and cancer (Mancini & Toker, 2009). For example, proliferation and anchorage-independent growth of pancreatic tumor cells is dependent on calcineurin and NFATc1 activity, consistent with the high levels of nuclear NFATc1 found in pancreatic tumors (Singh *et al.*, 2010). Likewise, our results found that VEGF-A was a potent inducer of NFATc1 expression, which may be required for epithelial cell migration and tubulogenesis. Although specific NFATc isoforms were not distinguished in the model, our simulations suggested that NFATc transcriptional activity was capable of maintaining epithelial traits, even during TGF-*β* induced EMT. Experimentally, we found that E-cadherin expression was dependent upon NFATc dephosphorylation in response to simultaneous VEGF-A and TGF-*β*1/2 treatment. Thus, these results support the hypothesis that NFATc activity plays a critical role in maintaining cell-cell contacts, even during partial EMT.

Epithelial cells reproduce tissue-like organization when grown in a three-dimensional extracellular matrix (ECM) environment, and therefore are an attractive model to study morphogenic mechanisms. It is well established that MCF10A cells form structures that closely resemble acini (multi-lobed cluster of cells) in three-dimensional in vitro cultures (Debnath *et al.*, 2003). It has been postulated that a cellular response reminiscent of partial EMT underlies this process, stimulating further branching and formation of acini (Pearson & Hunter, 2007). Normally well controlled process such as tubulogenesis can be co-opted by cancer cells to break away from a primary lesion and invade through the surrounding stroma (O’Brien *et al.*, 2004). However, by retaining a transient hybrid EMT-like state, clusters of these tube-forming tumor cells can reform at a high rate after invasion, possibly explaining why invasive human carcinomas frequently appear to be cellular collections with varying degrees of gland-like differentiation (Debnath & Brugge, 2005). In this study, we showed that our predicted hybrid phenotype generated by simultaneous treatment of epithelial cells with VEGF-A and TGF-*β* possessed altered migration and invasion, which enhanced tubular branching. A salient feature of this behavior, however, was the retention of cell-cell contacts that allowed cells to migrate without completely dissociating from their neighbors. Thus, our results support a mechanism in which hybrid cells can maintain some functional characteristics of epithelial cells such as cell-cell adhesion, which are normally lost in a fully differentiated mesenchymal state. The tumor microenvironment contains many soluble signals simultaneously, including VEGF and TGF-*β.* Thus, it is likely that some cancerous epithelial cells could exhibit hybrid EMT phenotypic states. This may explain why fibroblastoid morphology, a classical feature of EMT, is not commonly observed in human carcinomas (Debnath & Brugge, 2005). This study focused on the combinatorial effects of two very different ligand families present together in the tumor environment. Additional modeling studies are required to unravel the global response of epithelial cells to the full spectrum of chemical, substrate, and mechanical cues. The simulation strategy presented here is readily adaptable to larger species sets, with the major advantage that experimentally testable hypotheses can be generated regarding how signals get integrated to produce global cellular response. Furthermore, by simulating multiple ensembles of parameter sets, subpopulations across a constellation of phenotypes can be created and mined for common and/or divergent signaling characteristics. This is a significant advantage over forced convergence to a single unique solution and thereby generating a potentially non-physiological homogeneous population.

The deterministic population of EMT models predicted heterogeneous behavior that was qualitatively consistent with experimental studies. There is a diversity of algorithmic approaches to estimate model parameters (Moles *et al.*, 2003), as well as many strategies to integrate model identification with experimental design (Rodriguez-Fernandez *et al.*, 2013, Villaverde & Banga, 2014). However, despite these advances, the identification of models describing intracellular network behavior remains challenging. There are different schools of thought to deal with this challenge. One school has focused on model reduction. Data-driven approaches (Cirit & Haugh, 2012), boolean (Choi *et al.*, 2012) or other logical model formulations (Morris *et al.*, 2011, Terfve *et al.*, 2012) are emerging paradigms that constrain model complexity by the availability of the training and validation data. Other techniques such as constraints based modeling, which is commonly used to model metabolic networks, have also been applied to model transcriptional networks, although primarily in lower eukaryotes and prokaryotes (Hyduke & Palsson, 2010). These techniques (and many others, see review (Wayman & Varner, 2013)) are certainly exciting, with many interesting properties. Here, we used a traditional approach of mass action kinetics within an ordinary differential equation framework that also included transfer functions to simplify scenarios where reactions involving one species are controlled by several others (e.g. E-cadherin transcription). The identification problem for the EMT model was underdetermined (not uncommon for differential equation based models). However, a central criticism leveled by biologists is that model simplification is often done at the cost of biological reality, or done for reasons of computational expediency (Sainani, 2012). To avoid this criticism, we systematically identified an ensemble of likely models each consistent with the training data, instead of a single but uncertain best fit model. Previously, we (and others) have suggested that deterministic ensembles could model heterogeneous populations in situations where stochastic computation was not feasible (Lequieu *et al.*, 2011). Population heterogeneity using deterministic model families has previously been explored for bacterial growth in batch cultures (Lee *et al.*, 2009). In that case, distributions were generated because the model parameters varied over the ensemble, i.e., extrinsic noise led to population heterogeneity. In this study, parameters controlling physical interactions such as disassociation rates, or processes such as gene expression were distributed over the ensemble. Population heterogeneity can also arise from intrinsic thermal fluctuations, which are not captured by a deterministic population of models (Swain *et al.*, 2002). Thus, deterministic ensembles, provide a coarse-grained or extrinsic-only ability to simulate population diversity. Despite this limitation, our prediction of phenotypic heterogeneity (and the underlying signaling events responsible for the heterogeneity) was consistent with experimental observations. This suggested that deterministic ensembles could simulate disease or developmental processes in which heterogeneity plays an important role, without having to resort to stochastic simulation.

A common criticism of ODE modeling has been the poorly characterized effect of structural and parametric uncertainty. In this study, parametric uncertainty was addressed by developing an ensemble of probable models instead of a single best-fit but uncertain model using multiobjective optimization. While computationally complex, multiobjective optimization is an important tool to address qualitative conflicts in training data that arise from experimental error or cell line artifacts (Handl *et al.*, 2007). On the other hand, structural uncertainty is defined as uncertainty in the biological connectivity. The EMT model connectivity was assembled from an extensive literature review. However, several potentially important signaling mechanisms were not included. First, we identified a potential gap in biological knowledge surrounding the regulation of LEF-1 expression, that was filled by the addition of the hypothetical YREG1 transcriptional repressor. The LEF-1 transcription factor is expressed in tissues that undergo EMT during embryogenesis (Nawshad & Hay, 2003, Vega *et al.*, 2004), and has been suggested to promote an invasive phenotype in cancer cells (Cano *et al.*, 2000, Kim *et al.*, 2002). Low levels of YREG1 were important for stabilizing the interaction between LEF-1 and β-catenin, while elevated levels inhibited EMT by downregulating LEF-1 transcriptional activity. Recent evidence has established a complex role of Amino terminal Enhancer of Split (AES) and Groucho/TLE on suppressing LEF-1 activity. AES opposes LEF-1 transcriptional activation while Groucho/TLE binds with LEF-1 for a histone deacetylase repression. In addition, *β*-catenin directly displaces Groucho/TLE repressors from TCF/LEF-1 in Wnt-mediated transcription activation (Arce *et al.*, 2009, Grumolato *et al.*, 2013). Our model agrees with this newly discovered feedback system, as YREG1 regulates LEF-1 activity leading to EMT stabilization.

NF-κB may also play an essential role of in the epithelial transformation. NF-κB may influence Snail expression through the AKT pathway and directly stabilize Snail activity (Wu *et al.*, 2009). This is particularly important for integrating inflammation pathways, such as interleukin-6 (IL-6) and tumor necrosis factor-α (TNF-α), which have been linked to EMT in pathological conditions (Sullivan *et al.*, 2009). Other pathways such as Notch have also been shown to act synergistically with TGF-*β* to express Slug in the developing embryo (Niessen *et al.*, 2008). Lastly, while we have modeled classical protein signaling, we have not considered the role of regulatory RNAs on EMT. There is growing evidence that microRNAs (miRNAs) play a strong role in EMT, where several miRNAs, for example miR-21 and miR-31 are strongly associated with TGF-*β* exposure (Bullock *et al.*, 2012). Addressing missing structural components like these, could generate more insight into TGF-*β* signaling and its role in phenotypic transformation.

## Materials and Methods

The model code and parameter ensemble is freely available under an MIT software license and can be downloaded from http://www.varnerlab.org.

### Signaling network connectivity

The EMT model described the gene expression program resulting from TGF-*β* and VEGF-A signaling in a prototypical epithelial cell. The TGF-*β*-EMT network contained 97 nodes (proteins, mRNA or genes) interconnected by 251 interactions. The network connectivity was curated from more than 40 primary literature sources in combination with on-line databases (Jensen *et al.*, 2009, Linding *et al.*, 2007). The model interactome was not specific to a single epithelial cell line. Rather, we assembled canonical pathways involved in TGF-*β* and VEGF-A signaling, defaulting to human connectivity when possible. Using a canonical architecture allowed us to explore general features of TGF-*β* induced EMT without cell line specific artifacts.

Our signaling network reconstruction was based on Medici *et al.* who identified the pathways though which MDCKII, DLD1 colon carcinoma, and A375 melanoma cells transition towards a mesenchymal phenotype (Medici *et al.*, 2008). Sequential activation of MAPK and Smad pathways were initiated upon addition of TGF-*β*1/2. Briefly, TGF-*β*2 signals through the RAS-RAF-MEK-ERK pathway to up-regulate Snail and Slug expression (Medici *et al.*, 2006). Snail, a known repressor of junctional proteins, inhibits the expression of E-cadherin (Cano *et al.*, 2000). This initial repression of E-cadherin leads to a release of *β*-catenin from the cell membrane. This release of *β*-catenin can then translocate to the nucleus and form transcriptional complexes with TCF-4 to drive TGF-*β*3 expression (Medici *et al.*, 2008). The PI3K to GSK3 pathway was included and acted as an activating mechanism of *β*-catenin signaling through TGF-*β*3 signaling (Medici *et al.*, 2008). GSK3 is known to act on *β*-catenin signaling through the ubiquitin-proteasome pathway (Larue & Bellacosa, 2005, Zhou *et al.*, 2004). Thereby, further *β*-catenin release also resulted from by TGF-*β*3 signals to the cells interior by binding to type II receptors, which form heterodimers with type I receptors (ALK5) (Derynck & Zhang, 2003). This activates the receptors serine/threonine kinase activity to phosphorylate and activate the receptor Smads 2/3 (Massagué *et al.*, 2005). In the model, receptors are simplified and represented as either bound or unbound complexes with their ligands. Phosphorylated Smads 2/3 (pSmad2/3) form heterodimers and translocate to the nucleus. pSmads complexes up-regulate other transcription factors, such as LEF-1. The pSmad2/4-LEF-1 complex has been shown to directly repress the E-cadherin gene (Nawshad *et al.*, 2007). LEF-1 also binds with *β*-catenin to upregulate mesenchymal proteins such as fibronectin (Medici *et al.*, 2011). In the model, Smad signaling is consolidated into a single Smad species that can act in a co-dependent fashion with LEF1 to downregulate E-cadherin via a transfer function, eliminating the need for an explicity LEF-1, pSmad complex. The EMT gene expression program was initiated by the binding of TGF-*β* isoforms to TGF-*β* surface receptors, starting the downstream signaling program. Repression of E-cadherin expression is the central event in the transition from an epithelial to a mesenchymal phenotype (Cano *et al.*, 2000).However, this transition is not solely driven by transcriptional events. At the protein level, the repression of E-cadherin leads to a release of *β*-catenin from cell membrane. Cystolic *β*-catenin then translocates to the nucleus and forms transcriptionally-active complexes with immunoglobulin transcription factor 2 (TCF-4) to drive TGF-*β*3 expression (Medici *et al.*, 2008). The PI3K to GSK3 pathway was included and acted as an activating mechanism of *β*-catenin signaling through TGF-*β*3 signaling (Medici *et al.*, 2008). GSK3 is known to act on *β*-catenin signaling through APC complex associated ubiquitin-proteasome pathway. The APC complex is represented in our model and serves as a second reservoir of *β*-cateinin in untransformed cells whose sequestration is regulated by GSK3 (Larue & Bellacosa, 2005, Medici *et al.*, 2008, Zhou *et al.*, 2004). Lastly, VEGF-A activation of NFATc1 takes place through calcineurin signaling leading to an enhancement of E-cadherin expression (Suehiro *et al.*, 2014), as supported by our VEGF-A experimental data (Fig. S4).

### Formulation, solution and analysis of the EMT model equations

#### EMT signaling events

EMT signaling events were modeled using either saturation or mass-action kinetics within an ordinary differential equation (ODE) framework:

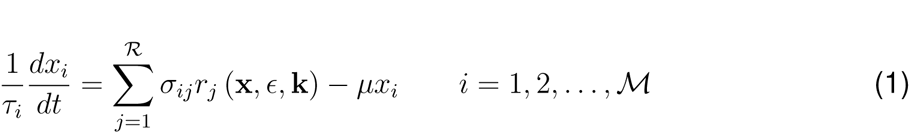

where 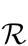 denotes the number of signaling reactions and 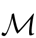 denotes the number of signaling proteins in the model. The quantity *τ*_*i*_ denotes a time scale parameter for species *i* which captures un-modeled effects; in the current study *τ*_*i*_ = 1 for all species. The quantity *r*_*j*_(x,ϵ,k) denotes the rate of reaction *j.* Typically, reaction *j* is a non-linear function of biochemical and enzyme species abundance, as well as unknown model parameters k 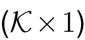. The quantity *σ*_*ij*_ denotes the stoichiometric coefficient for species *i* in reaction *j.* If *σ_*ij*_ >*0, species *i* is produced by reaction *j.* Conversely, if *σ*_*ij*_ < *0*, species *i* is consumed by reaction *j*, while *σ*_*ij*_ = 0 indicates species *i* is not connected with reaction *j.* Species balances were subject to the initial conditions x (*t*_*o*_) = x_*o*_.

Rate processes were written as the product of a kinetic term 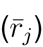 and a control term (*v*_*j*_) in the EMT model. The rate of enzyme catalyzed reactions was modeled using saturation kinetics:

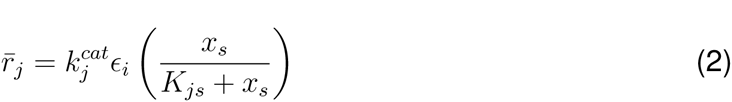

where 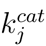 denotes the catalytic rate constant for reaction *j*, *ε*_*i*_ denotes the abundance of the enzyme catalyzing reaction *j*, and *K*_*js*_ denotes the saturation constant for species *s* in reaction *j.* On the other hand, mass action kinetics were used to model protein-protein binding interactions within the network:

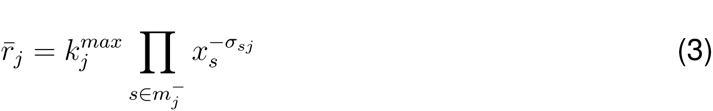

where 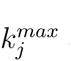 denotes the maximum rate for reaction *j*, *σ*_*sj*_ denotes the stoichiometric coefficient for species *s* in reaction *j*, and *s ϵ m*_*j*_ denotes the set of *reactants* for reaction *j.*

Reversible binding was decomposed into two irreversible steps.

The control terms 0 ≤ *v*_*j*_ ≤ 1 depended upon the combination of factors which influenced rate process *j.* For each rate, we used a rule-based approach to select from competing control factors. If rate *j* was influenced by 1,…, *m* factors, we modeled this relationship as 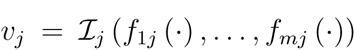 where 0 ≤ *f*_*ij*_(·) ≤ 1 denotes a regulatory transfer function quantifying the influence of factor *i* on rate *j.* The function 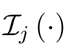 is an integration rule which maps the output of regulatory transfer functions into a control variable. In this study, we used 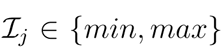 and hill transfer functions (Sagar & Varner, 2015, Wayman *et al.*, 2015). If a process has no modifying factors, *v*_*j*_ = 1.

#### EMT gene expression processes

The EMT model described both signal transduction and gene expression events following the addition of TGF-*β* and VEGF-A. For each gene of the 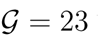 we considered, we modeled both the resulting mRNA (*m*_*j*_) and protein (*p*_*j*_):

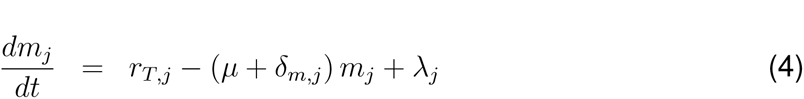

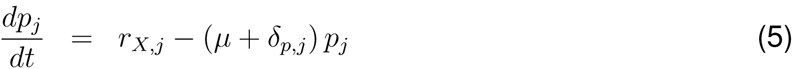

where 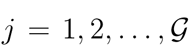. The terms *rT*,_*j*_ and *rX*,_*j*_ denote the specific rates of transcription, and translation while the terms *δ*_*m,j*_ and *δ*_*p,j*_ denote degradation constants for mRNA and protein, respectively. Lastly, *μ* denotes the specific growth rate, and *λ*_*j*_ denotes the constitutive rate of gene expression for gene *j.* The specific transcription rate was modeled as the product of a kinetic term 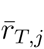 and a control term *u*_*j*_ which described how the abundance of transcription factors, or other regulators influenced the expression of gene *j.* The kinetic rate of transcription was modeled as:

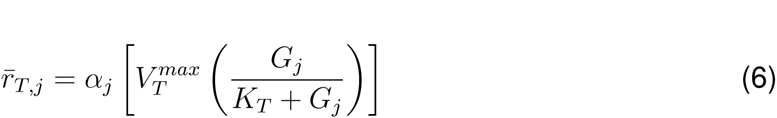

where the maximum gene expression rate was defined as the product of a characteristic transcription rate constant (*k*_*T*_) and the abundance of RNA polymerase, 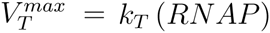. The parameter *α*_*j*_ denotes the gene specific correction to the characteristic transcription rate. (estimated in this study), while *k*_*T*_, *G*_*j*_ and RNAP were estimated from literature (Milo *et al.*, 2010). Similar to the signaling processes, the gene expression control term 0 ≤ *u*_*j*_ ≤ 1 depended upon the combination of factors which influenced rate process *j.* For each rate, we used a rule-based approach to select from competing control factors. If the expression of gene *j* was influenced by 1,…, *m* factors, we modeled this relationship as 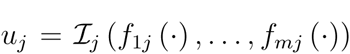 where 0 ≤ *f*_*ij*_ (·) ≤ 1 denotes a regulatory transfer function quantifying the influence of factor *i* on the expression of gene *j.* The function 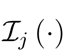 is an integration rule which maps the output of regulatory transfer functions into a control variable. In this study, we used 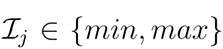 and hill transfer functions (Sagar & Varner, 2015, Wayman *et al.*, 2015). If a gene expression process has no modifying factors, *u*_*j*_ = 1.

Lastly, the specific translation rate was modeled as:

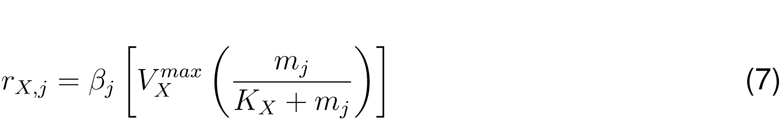

where 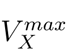 denotes a characteristic maximum translation rate estimated from literature, *β*_*j*_ denotes the transcript specific correction the characteristic translation rate, and *K*_*X*_ denotes a translation saturation constant. The characteristic maximum translation rate was defined as the product of a characteristic translation rate constant (*k*_*X*_) and the abundance of Ribosomes (*RIBO*), 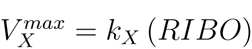, where both *k*_*X*_ and *RIBO* abundance were estimated from literature (Milo *et al.*, 2010).

In this study, we estimated *k*_*T*_,*k*_*X*_,*RNAP* and *RIBO* from literature. We calculated the concentration for gene *j* by assuming, on average, the cell had two copies of each gene at any given time. We also estimated characteristic values for *δ*_*m,j*_ and *δ*_*p,j*_, and corrected these values with mRNA/protein specific correction factors. Thus, the bulk of our gene expression parameters were based directly upon literature values. The values used for the characteristic transcription/translation parameters are given in the supplemental materials.

The signaling and gene expression model equations were implemented in Julia and solved using the CVODE routine of the Sundials package (Bezanson *et al.*, 2014, Hindmarsh *et al.*, 2005). The model code and parameter ensemble is freely available under an MIT software license and can be downloaded from http://www.varnerlab.org.

#### Estimation of model parameters using multiobjective optimization

We estimated a population of likely EMT model parameter sets (each consistent with the training data) using 41 data sets generated in DLD1 colon carcinoma, MDCKII, and A375 melanoma cells taken from Medici *et al.* (Medici *et al.*, 2008). We used the Pareto Optimal Ensemble Technique (JuPOETs) multiobjective optimization framework in combination with leave-one-out cross-validation to estimate an ensemble of model parameters (Bassen *et al.*, 2016, Song *et al.*, 2010). Model parameter values were adjusted to minimize the residual between simulations and experimental measurements for the 11 objective functions (supplemental materials). Cross-validation was used to calculate both training and prediction error during the parameter estimation procedure (Kohavi, 1995). The 41 intracellular protein and mRNA data-sets used for identification were organized into 11 objective functions. These 11 objective functions were then partitioned, where each partition contained ten training objectives and one validation objective. The training and validation data were Western blots. We achieved a biologically realistic concentration scale by establishing characteristic rates of transcription, translation, mRNA and protein degradation, as well as characteristic concentrations of ribosomes and RNAPs using the Bionumbers database (Milo *et al.*, 2010), (supplemental materials). The overall concentration scale was nM, with proteins ranging from 10-1000nM and mRNA ranging from 0.01 to 1nM, reflecting the true abundances and ratios between each species. An initial nominal parameter set was established by inspection. JuPOETs was then allowed to search in a neighborhood of ±30% of this nominal set. The parameter ensemble estimated by JuPOETs is available with the model source code. JuPOETs is open source and freely available for download under an MIT software license from http://www.varnerlab.org.

### Cell culture and experimental interrogation

DLD1 colon carcinoma, MCF10A, and HUVEC were acquired from the American Tissue Culture Collection (Manassas, VA). Cells were grown in culture with RPMI 1640 medium with 10% fetal bovine serum and 1% penicillin/streptomycin for DLD1, EBM-2 supplemented with EGM-2, 5% fetal bovine serum, and 1% penicillin/streptomycin for HUVEC, or MGEM 2 supplemented with insulin, bovine pituitary extract, cholera toxin, hEGF, hydrocortisone, 5% horse serum, and 1% penicillin/streptomycin for MCF10A. Cells were serum starved for 24 hours and removed from all experimental conditions. Recombinant VEGFA165 was also removed from culture medium prior to experimentation. Recombinant human TGF-*β*2 (R & D Systems, Minneapolis, MN) was added to the culture medium at a concentration of 10 ng/ml and recombinant VEGFA165 at a concentration of (5ng/ml, 50ng/ml) for all relative experiments. NFAT inhibitor (VIVIT peptide) (EMDBiosciences, Darmstadt, Germany), was added to the culture medium at a concentration of 10*μ*M for all relative experiments. Cells were passaged 1:3 or 1:4 every 3-6 d and used between passages 4 and 8.

#### VEGF treatment

DLD1 and MCF10A cells were suspended in culture media (with RPMI 1640 medium with 10% fetal bovine serum and 1% penicillin/streptomycin for DLD1 or MGEM 2 supplemented with insulin, bovine pituitary extract, cholera toxin, hEGF, hydrocortisone, 5% horse serum, and 1% penicillin/streptomycin for MCF10A), and allowed to aggregate overnight in hanging drop culture (20*μ*L; 20,000 cells). The spherical aggregates were placed on the surface of neutralized type I collagen hydrogels (1.5mg/mL) and allowed to adhere. Cultures were then serum starved (1% serum) for 24 hours. Recombinant VEGFA165 was then added to the media (5ng/ml, 50ng/ml) and mRNA was harvested after 3hr and 24hr timepoint.

#### RT-PCR

RNA extractions were performed using a Qiagen total RNA purification kit (Qiagen, Valencia, CA) and RNA was reverse transcribed to cDNA using the SuperScript III RT-PCR kit with oligo(dT) primer (Invitrogen). Sufficient quality RNA was determined by an absorbance ratio A260/A280 of 1.8-2.1, while the quantity of RNA was determined by measuring the absorbance at 260nm (A260). Real-time PCR experiments were conducted using the SYBR Green PCR system (Biorad, Hercules, CA) on a Biorad CFX96 cycler, with 40 cycles per sample. Cycling temperatures were as follows: denaturing, 95C; annealing, 60C; and extension, 70C. Primers were designed to detect GAPDH, E-cadherin, vimentin, Slug, Sp1, and NFATc1 in cDNA clones: Sp1 (F-TTG AAA AAG GAG TTG GTG GC, R-TGC TGG TTC TGT AAG TTG GG, Accession NG030361.1), NFATc1 (F-GCA TCA CAG GGA AGA CCG TGT C, R-GAA GTT CAA TGT CGG AGT TTC TGA G, Accession NG029226.1). GAPDH, E-cadherin, vimentin, and Slug primers were taken from previously published literature (Medici *et al.*, 2008).

#### Antibody Staining

Samples were fixed in 4% PFA overnight at 4C. Samples were then washed for 15 minutes on a rocker 3 times with PBS, permeabilized with 0.2% Triton-X 100 (VWR International, Radnor, PA) for 10 minutes, and washed another 3 times with PBS. Samples were incubated overnight at 4C in a 1% BSA (Rockland Immunochemicals, Inc., Gilbertsville, PA) blocking solution followed by another 4C overnight incubation with either rabbit anti-human E-cadherin 1:100 (Abcam, ab53033), mouse anti-human phospho-Sp1 1:100 (Abcam, ab37707), mouse anti-human vimentin 1:100 (Invitrogen, V9), and rabbit anti-human NFATc1 (Santa Cruz, sc-7294) 1:100. After 3 washes for 15 minutes with PBS, samples were exposed to Alexa Fluor 488 or 568 conjugated (Invitrogen), species specific secondary antibodies at 1:100 in 1% BSA for 2 hours at room temperature. Three more washes with PBS for 15 minutes were followed by incubation with either DRAQ5 far red nuclear stain (Enzo Life Sciences, Plymouth Meeting, PA) at 1:1000.

#### FACS

Flow cytometry for E-cadherin 1:100 (Abcam) and vimentin 1:100 expressing cells was performed. Briefly, cells were trypsinized, fixed with 4% PFA for 10 min and then preserved in 50% methanol/PBS. Cells were kept in the −20C until antibody staining was preformed. Samples were divided into multiple aliquots in order to stain the proteins separately and compensate for secondary antibody non-specific binding. Cells were incubated for 24 hrs at 4 C in primary antibody diluted in either PBS (extracellular) or 0.2% saponin-PBS (intracellular). Cells were then washed 3 times with PBS and incubated with appropriate secondary antibodies and imaged using a Coulter Epics XL-MCL Flow Cytometer (Coulter). All samples were compensated using appropriate background subtraction and all samples were normalized using 7500 cells per flow condition.

#### Three-Dimensional Culture and Tubulogenesis Assays

For invasion/migration assays, cells were resuspended in culture media, and allowed to aggregate overnight in hanging drop culture (20*μ*L; 20,000 cells). The spherical aggregates were placed on the surface of neutralized type I collagen hydrogels (1.5mg/mL) and allowed to adhere for 2 hrs before adding treatments. Cultures were maintained for 72 hrs, after which they were fixed in 4% PFA and slowly rehydrated using PBS. For compaction assays, cells were pelleted via centrifugation and resuspended within a neutralized collagen hydrogel (1.5mg/mL) solution at a density of 400,000 cells/mL. 250*μ*L of gel was inoculated into culture wells, which solidified after 60min. Treatments were then added within 800*μ*L of the culture medium without serum. Gels were liberated from the surfaces of the culture wells the next day and cultured free floating for an additional 3-7 days, exchanging serum free media with appropriate factors every 48 hrs.

Tubulogenesis was defined as a typical nonmalignant acini structure. This includes a polarized epithelial cell, hollow lumen, and the basal sides of the cell are surrounded by ECM proteins (Fig. 6A, Controls or VEGF treated). Previous work has shown that change in the morphological characteristics of nontumorigenic MCF10A epithelial acini occur over time and exploiting them to growth in 3D culture can be quantified. For example, using image segmentation, Chang *et al.* (Chang *et al.*, 2007) examined the elongation of the MCF10A acini at 6, 12, and 96 hours after a particular treatment. Polizzotti *et al.* (Polizzotti *et al.*, 2012) also suggested a computational method to quantify acini structure based on morphological characteristics in nonmalignant, noninvasive, and invasive conditions.

Adapted from these approaches, we first fluorescently labeled our cultures and captured the acini structures by 3D confocal microscopy. Next individual acini structures in the images were segmented by imageJ and labeled. We then extracted the number of ductal branches. Ductal branching was defined as any elongated cell cluster extending away from the total acini structure, which was manually segmented and counted using ImageJ. A total of 5 images for each condition were used, and approximately 12 acini were analyzed in each image. Total branching was normalized to the amount of acini present, and provides an overall general assessment to the extent of acini remodeling.

#### Statistics

Results are expressed as mean **±** standard error, n≥6. Data was analyzed with the GraphPad Prism version 4.00 for Windows (GraphPad Software, San Diego, CA) and SAS (Statistical Analysis Software, Cary, NC). A one-way ANOVA with Tukey's post hoc was used to compare differences between means and data was transformed when necessary to obtain equal sample variances. Differences between means were considered significant at p≺0.05.

## Supplemental Materials and Methods

### Characteristic transcription and translation parameters

We used literature based transcription and translation parameters to establish the characteristic synthesis and degradation rates for both mRNA and protein. We estimated values for the rate parameters from the Bionumbers database Milo *et al.* (2010). These parameters were then used for all gene expression calculations:

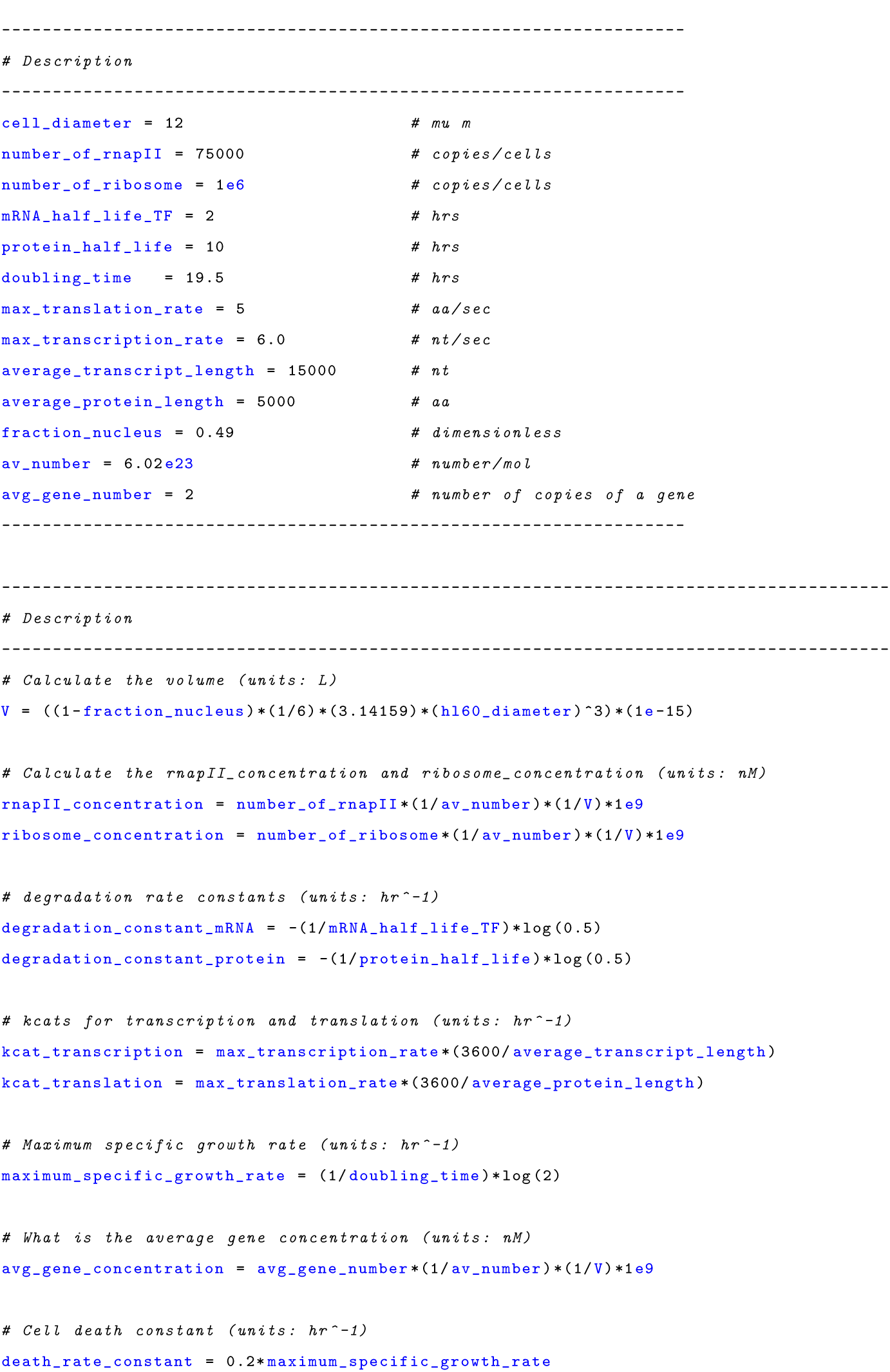
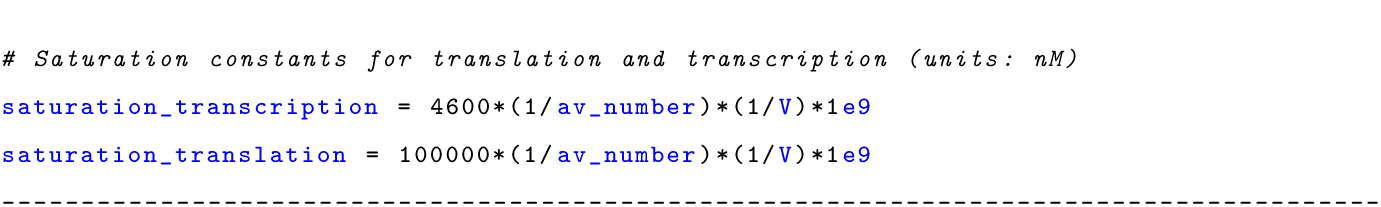

### Estimation and cross-validation of EMT model parameters

We used the Pareto Optimal Ensemble Technique (POETs) multiobjective optimization framework in combination with leave-one-out cross-validation to estimate an ensemble of TGF-*β*/EMT models. Cross-validation was used to calculate both training and prediction error during the parameter estimation procedure Kohavi (1995). The 41 intracellular protein and mRNA data-sets used for identification were organized into 11 objective functions. These 11 objective functions were then partitioned, where each partition contained ten training objectives and one validation objective. POETs integrates standard search strategies e.g., Simulated Annealing (SA) or Pattern Search (PS) with a Pareto-rank fitness assignment Bassen *et al.* (2016), Song *et al.* (2010). Denote a candidate parameter set at iteration *i* + 1 as k_*i*+1_. The squared error for k_*i*+1_ for training set *j* was defined as:

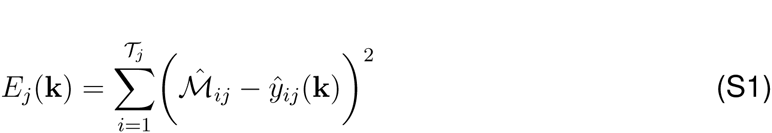

The symbol 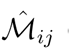 denotes scaled experimental observations (from training set *j*) while 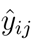 denotes the scaled simulation output (from training set *j*). The quantity *i* denotes the sampled time-index and 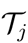 denotes the number of time points for experiment *j.* In this study, the experimental data used for model training was typically the band intensity from Western or Northern blots. Band intensity was estimated using the ImageJ software package Abramoff *et al.* (2004). The scaled measurement for species *x* at time *i* = {*t*_1_,*t*_*2*_, ..,*t*_*n*_} in condition *j* is given by:

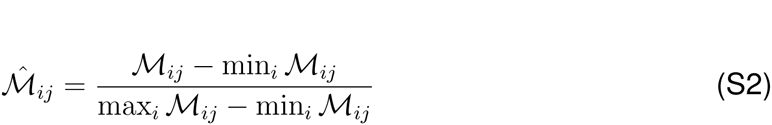

**Fig. S1:**
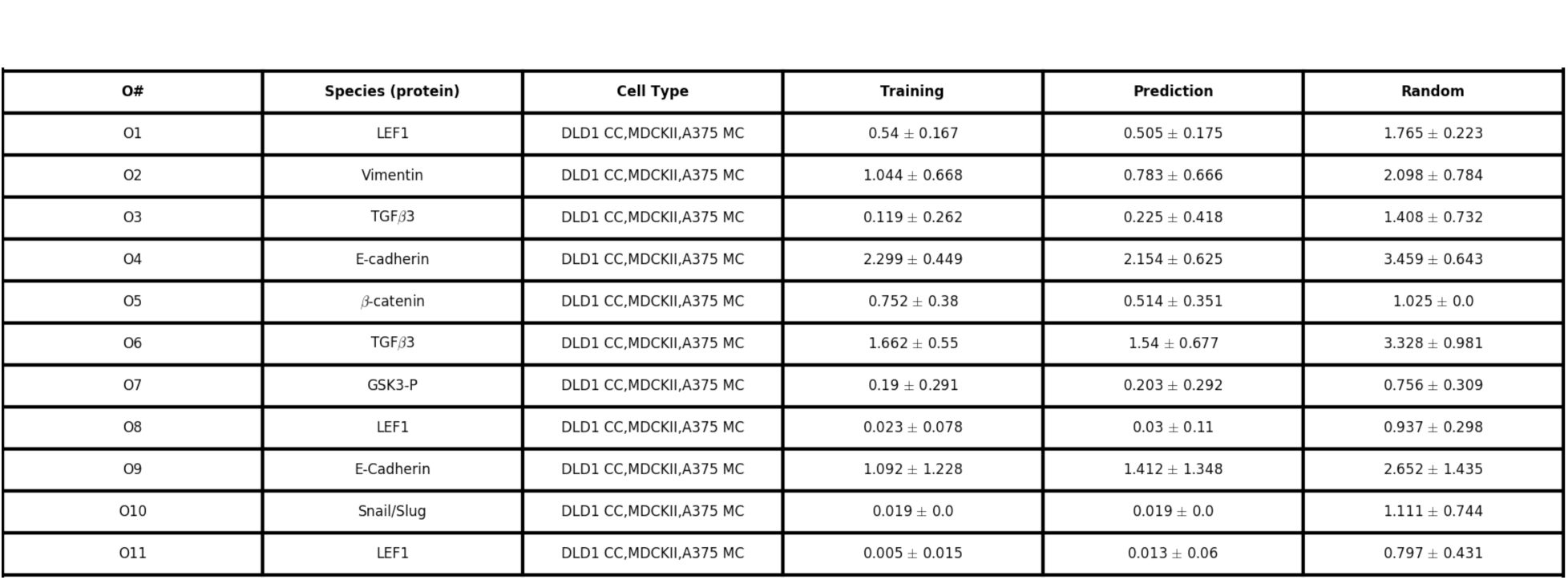
Training and prediction values as a function of condition for the 11 TGF-*β* objective functions versus a random parameter control.

Under this scaling, the lowest intensity band equaled zero while the highest intensity band equaled one. A similar scaling was defined for the simulation output. By doing this scaling, we trained the model on the relative change in blot intensity, over conditions or time (depending upon the experiment). Thus, when using multiple data sets (possibly from different sources) that were qualitatively similar but quantitatively different e.g., slightly different blot intensities over time or condition, we captured the underlying trends in the scaled data. JuPOETs is free or charge, open source and available for download under an MIT software license from http://www.varnerlab.org. Details of the JuPOETs implementation, including example codes are presented in Bassen *et al.*, Bassen *et al.* (2016).

**Fig. S2:**
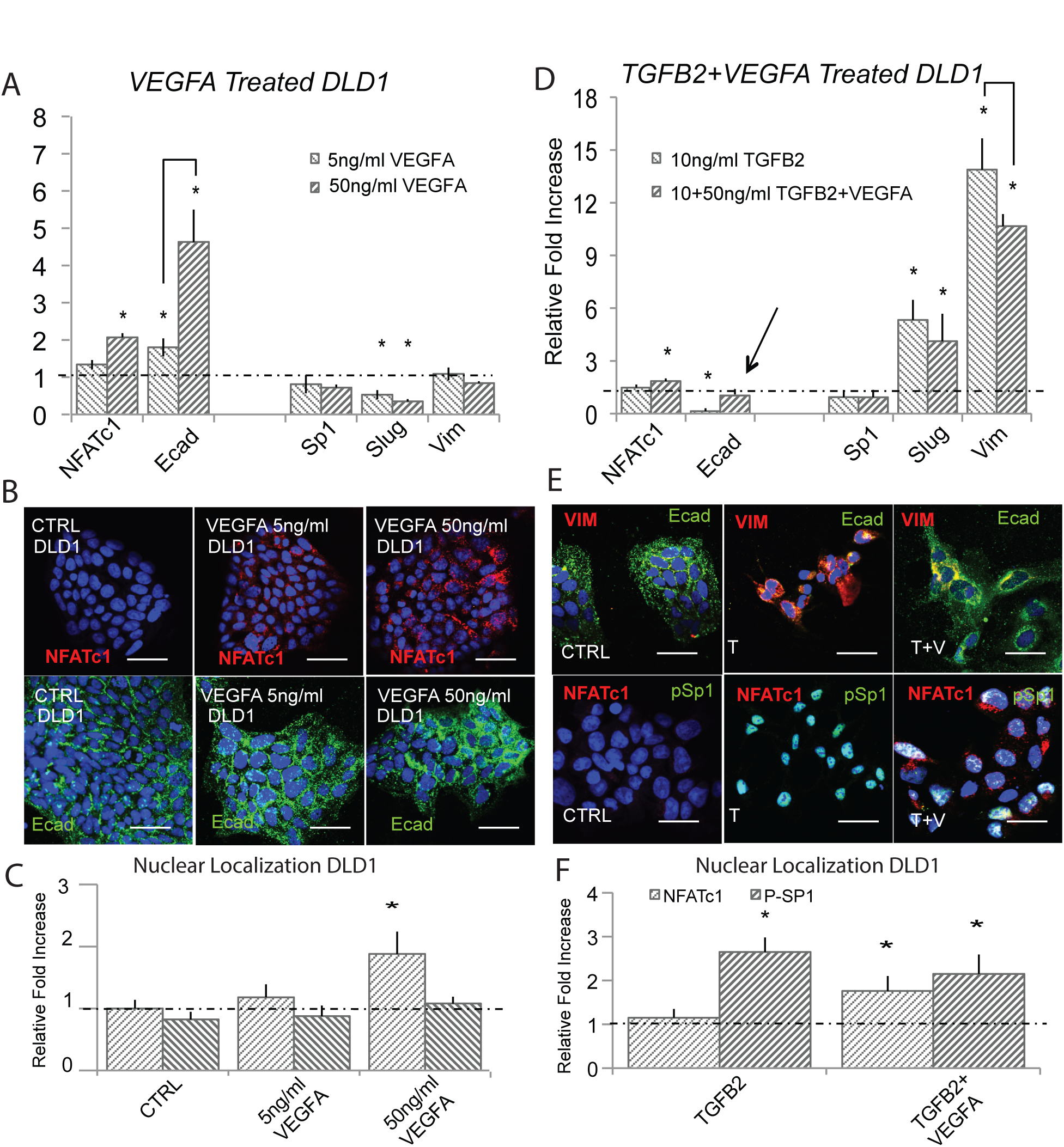
VEGF-A attenuates TGF-*β*1/2 to induce phenotype heterogeneity in DLD1. (A) In DLD1, we found that 5ng/ml of VEGFA increased NFATc1 and E-cadherin gene expression via qPCR and 50ng/ml potentiated this effect at 48 hrs. (B - C) These findings were confirmed at the protein level via immunofluorescence, as ecadherin levels and nuclear localization of NFATc1 increased. (D) Treatment with (10ng/ml) TGF*β*2 resulted in mesenchymal transformation as measured via qPCR against target genes Slug, ecadherin, vimentin, Sp1, and NFATc1. (E - F) Immunofluorescence and nuclear localization revealed a strong presence of phospho-Sp1. (G) Combination of VEGFA (50ng/ml) and TGF*β*2 (10ng/ml) treatment resulted in increased Slug, NFATc1, and vimentin expression, while also increasing ecadherin levels compared to control. (H) Immunofluorescence confirmed these results, as both ecadherin and vimentin levels were elevated. (I) A significant increase in nuclear localization of both NFATc1 and phospho-Sp1 were also found. Magnification, 40x. Scale bars: 50*μ*m. C=Control, T=TGF*β*2, V=VEGFA, VI=NFAT inhibitor (VIVIT). Asterisks signify statistical differences from each other according to a one-way ANOVA with Tukey's post hoc (p≺0.05).

**Fig. S3:**
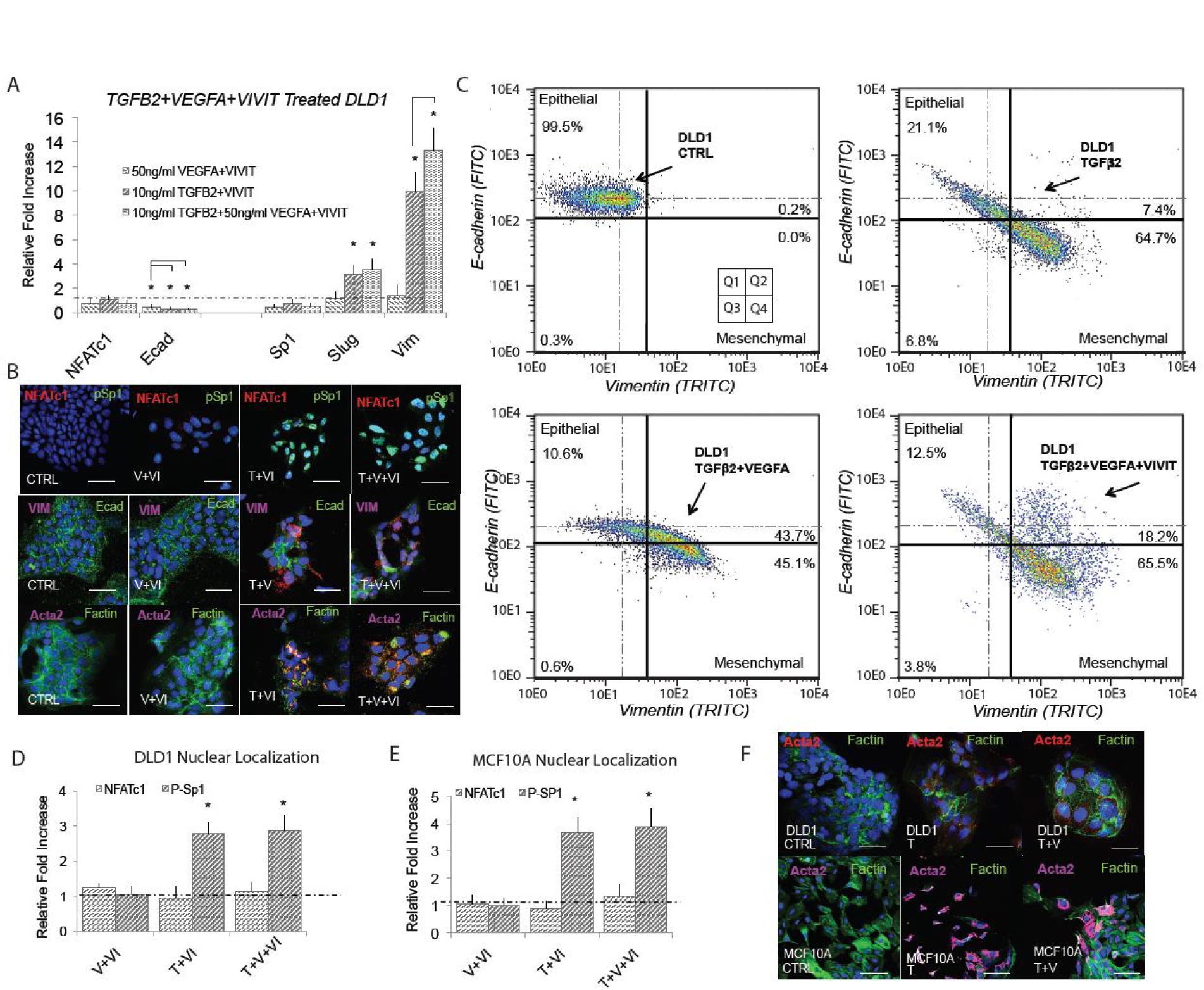
E-cadherin expression is dependent upon NFAT activity in DLD1. (A) Treatment with VEGFA (50ng/ml) and NFAT inhibitory peptide VIVIT (10*μ*M) resulted in significantly reduced ecadherin expression (qRT-PCR at 48hrs). Addition of TGF*β*2 (10ng/ml) and VIVIT resulted in increased Slug and vimentin expression, while inhibiting ecadherin levels. Combined TGF*β*2, VEGFA, and VIVIT treatment resulted in target genes Slug and vimentin expression increased, while inhibiting ecadherin levels. No change in Sp1 or NFATc1 expression was found. (B) These findings were confirmed via immunofluorescence as the VIVIT inhibitors was shown to inhibit ecadherin levels in all three cases. We also found no change in gene or nuclear localization of NFATc1 in all three cases, while phospho-Sp1 was found to increase in both TGF*β* conditions. (C) Quantitative flow cytometry also confirmed this trend. (D,E) TGF*β*2, VEGFA and VIVIT treatment in DLD1 and MCF10A resulted in no change of Sp1 expression or NFATc1 expression. (F) Likewise, no change in nuclear localization of NFAT in all three cases, however phospho-Sp1 was found to increase in both TGF*β* conditions. Magnification, 40x. Scale bars: 50*μ*m. C=Control, T=TGF*β*2, V=VEGFA, VI= NFAT inhibitor (VIVIT). Asterisks signify statistical differences from each other according to a one-way ANOVA with Tukey’s post hoc (p*≺*0.05).

**Fig. S4:**
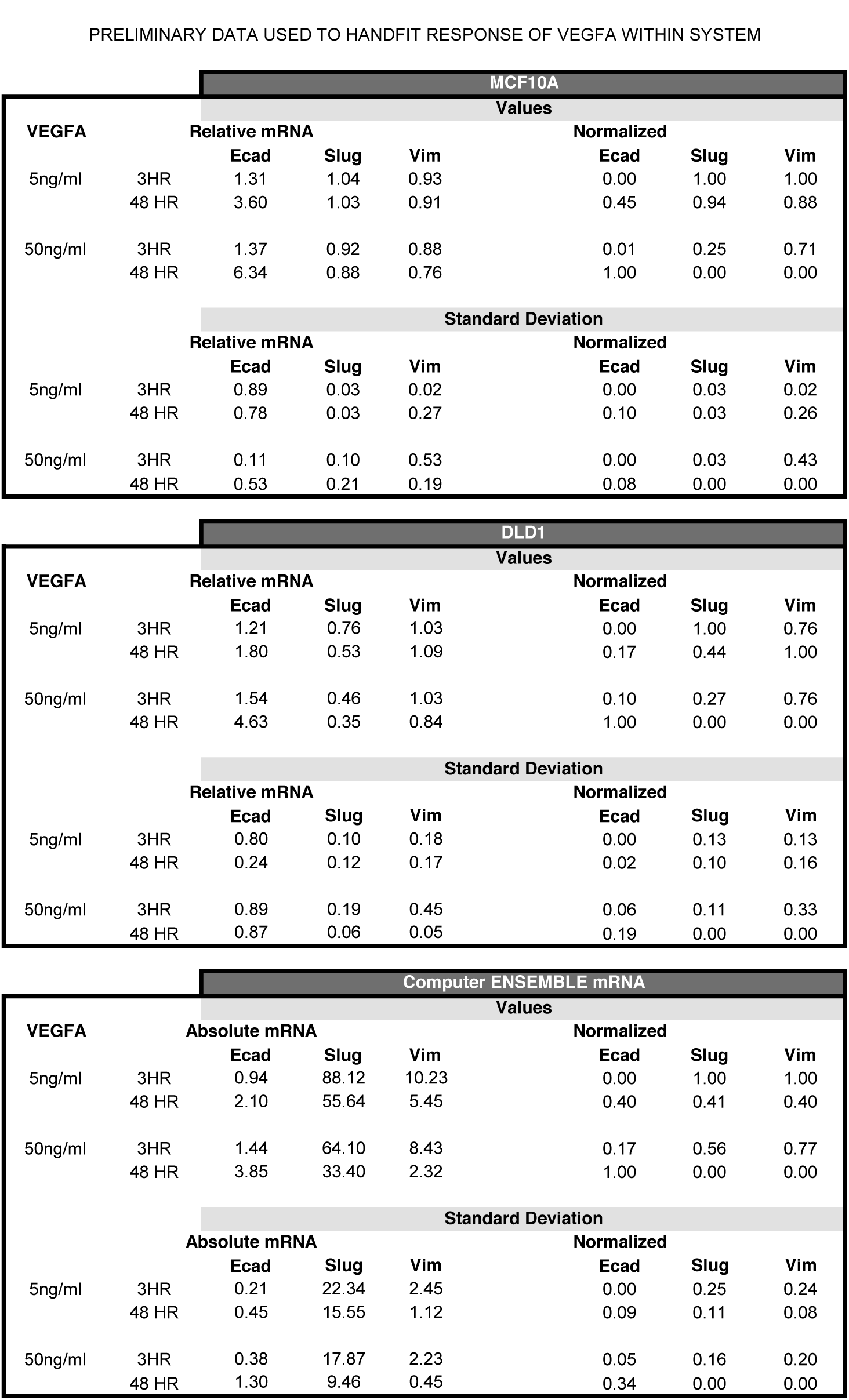
VEGF-A qPCR data used to hand fit VEGF enhancement of E-cadherin expression. mRNA was harvested after 3hr and 24hr timepoint.

